# A drought stress-induced MYB transcription factor regulates pavement cell shape in leaves of European aspen (*Populus tremula*)

**DOI:** 10.64898/2026.01.16.699252

**Authors:** Sijia Liu, Siamsa M. Doyle, Kathryn M. Robinson, Zahra Rahneshan, Nathaniel R. Street, Stéphanie Robert

## Abstract

The leaf pavement cells of many plant species develop fascinating jigsaw puzzle-like shapes in which neighboring cells interdigitate with each other, providing an ideal model for the study of cell shape acquisition. We analyzed pavement cell shape complexity in a natural population of European aspen (*Populus tremula*) genotypes and then used a genome-wide association study (GWAS) approach to identify a new candidate gene in cell shape regulation, Potra2n8c18226, encoding the transcription factor MYB305a. We subsequently validated a role for *MYB305a* in regulating aspen leaf pavement cell shape. We then demonstrated that drought conditions strongly induced *MYB305a* promoter expression in these cells and provided evidence implying that *MYB305a* plays a role in simplification of the cell shape in response to drought stress. Finally, we demonstrated negative correlations of pavement cell shape complexity with water-use efficiency, as well as with average precipitation, latitude and longitude of the genotypes’ original sampling sites in the natural European aspen population. Taken together, our results suggest that climatic variables affect the shape complexity of pavement cells in aspen leaf and provide a first step towards unravelling the molecular mechanisms controlling pavement cell shape acquisition in this species in response to environmental conditions, by implicating the involvement of the transcription factor MY305a.

## Introduction

In many plant species, the pavement cells of the leaf epidermis develop an intriguing jigsaw puzzle-like shape in which neighboring cells interdigitate with each other in a coordinated manner. Among several hypothesized functional roles for this cell shape, substantial computational modelling efforts support the idea of a role in alleviating the mechanical stress predicted to be generated in the epidermis of broad leaves during their near-isotropic expansion^1–3^. Interdigitated jigsaw puzzle-shaped pavement cells can be thought of as model cells for improving our understanding of how plants coordinate cell morphology in a tissue context. The cell wall ultimately defines the shape of plant cells. Studies of pavement cells, primarily in Arabidopsis (*Arabidopsis thaliana*) but also in a number of other species, have led to insights into how the cell wall determines this as well as the hormonal and molecular signaling pathways involved (for details, see suggested reviews^4–8^). Although *Populus* species also display jigsaw puzzle-shaped leaf pavement cells^6^, there has been little work done to investigate epidermal cell shape regulation in the *Populus* genus. These species are models for forest tree research due to their relatively small genome size, genetic diversity, amenability to transformation, ease of vegetative propagation, fast growth rate, usefulness for industrial production of paper materials, and potential for bioenergy production^9–11^. A fundamental understanding of growth and development in *Populus* is therefore important for the continued progression of biotechnological advances in these woody perennial species. We aimed therefore to investigate the regulation of leaf pavement cell shape in *Populus* species. On both the adaxial (upper) and abaxial (lower) sides of Arabidopsis leaves, the multiple asymmetrical divisions of small, undifferentiated meristemoid cells in stomatal cell lineage complexes is a major source of pavement cell origin^6,12^. The resulting young pavement cells subsequently develop outgrowths and interdigitate with their neighbors as they expand. In contrast, only the abaxial leaf side of European aspen (*Populus tremula*) develops stomata and the pavement cells on both leaf sides seem to arise from the rather symmetrical divisions of pre-existing interdigitated cells, resulting in straight cell walls that subsequently undulate as the sister cells continue to grow^6^. These rather dissimilar processes of cell development suggest that pavement cell shape regulatory mechanisms may be different between Arabidopsis and aspen. Furthermore, high diversity of pavement cell shape with shallow phylogenetic signal (low similarity among closely related species) were found in the eudicots^13^, further supporting the idea that eudicot species may have evolved different regulatory mechanisms for this process. This inspired us to seek novel regulators of pavement cell shape in aspen leaves. We analyzed leaf pavement cell shape complexity and size features in a natural population of European aspen genotypes, revealing variation in cell shape and size across the population. We then used a genome-wide association study (GWAS) approach to identify a new candidate gene, encoding a MYB family transcription factor, in cell shape regulation. We subsequently validated a role for the GWAS candidate, *MYB305a* (Potra2n8c18226), in aspen leaf pavement cell shape regulation through transgenic approaches. Since many MYB transcription factors play roles in drought stress responses^14^, we next applied drought treatments to our trees. We demonstrated that *MYB305a* promoter expression is strongly drought-induced in aspen leaf epidermis and found evidence supporting the involvement of *MYB305a* in regulating pavement cell shape responses to drought stress in this species.

## Results

### Variation in complexity of leaf pavement cell shape among European aspen genotypes in the field

To investigate the variability of leaf pavement cell shape complexity within one tree species, we took advantage of the Swedish aspen (SwAsp) collection of over 100 genotypes of European aspen that were collected from throughout Sweden and planted in a common garden near Umeå, Sweden^15^. We harvested leaves from the adult trees (so-called pre-formed leaves, since they emerge from buds each Spring) during the summer months, performed microscopy imaging of the adaxial leaf blade surface, and analyzed several shape- and size-related features of the pavement cells. The shape-related traits included circularity (measure of deviation from a perfect circle), branch count (number of branches in the cell skeleton), lobe number (number of cell outgrowths), solidity (ratio of cell area to area of the cell’s convex hull, a convex polygon fitting around the cell) and lobeyness (ratio of cell perimeter to convex hull perimeter)^1,16^. The size-related traits included non-lobe area (cell area excluding lobes), cell area, perimeter, length and width (lengths of the major and minor axes, respectively, of an ellipse fitting around the cell)^16^. Non-lobe area and perimeter might also be considered as being related to cell shape. We observed that these traits varied among the SwAsp genotypes (Supplementary Data S1). We then focused our attention on circularity, branch count, lobe number and area, and found clear variation in these features among the genotypes (Fig. 1A-D). We next selected four representative genotypes each for more and less complex pavement cell shape (indicated in magenta and cyan, respectively, in Fig. 1A-D and with the same color coding in subsequent figures) for later studies (shown in more detail in Supplementary Fig. 1A-D and Supplementary Fig. 2A-H). We also noted that pavement cell shape was generally not highly complex, even in the genotypes with more complex cell shape (Fig. S2A-H). In some cases, the degree of cell shape complexity appeared to be associated with cell area, since many genotypes with simple cell shapes had small cells, and vice versa (Fig. 1A-D). However, this was not always the case. For example, the genotypes SwAsp057 and 073 displayed larger cells with simpler cell shapes, while SwAsp078 and 088 displayed smaller cells with more complex cell shapes (Fig. 1A-D).

**Fig. 1:**
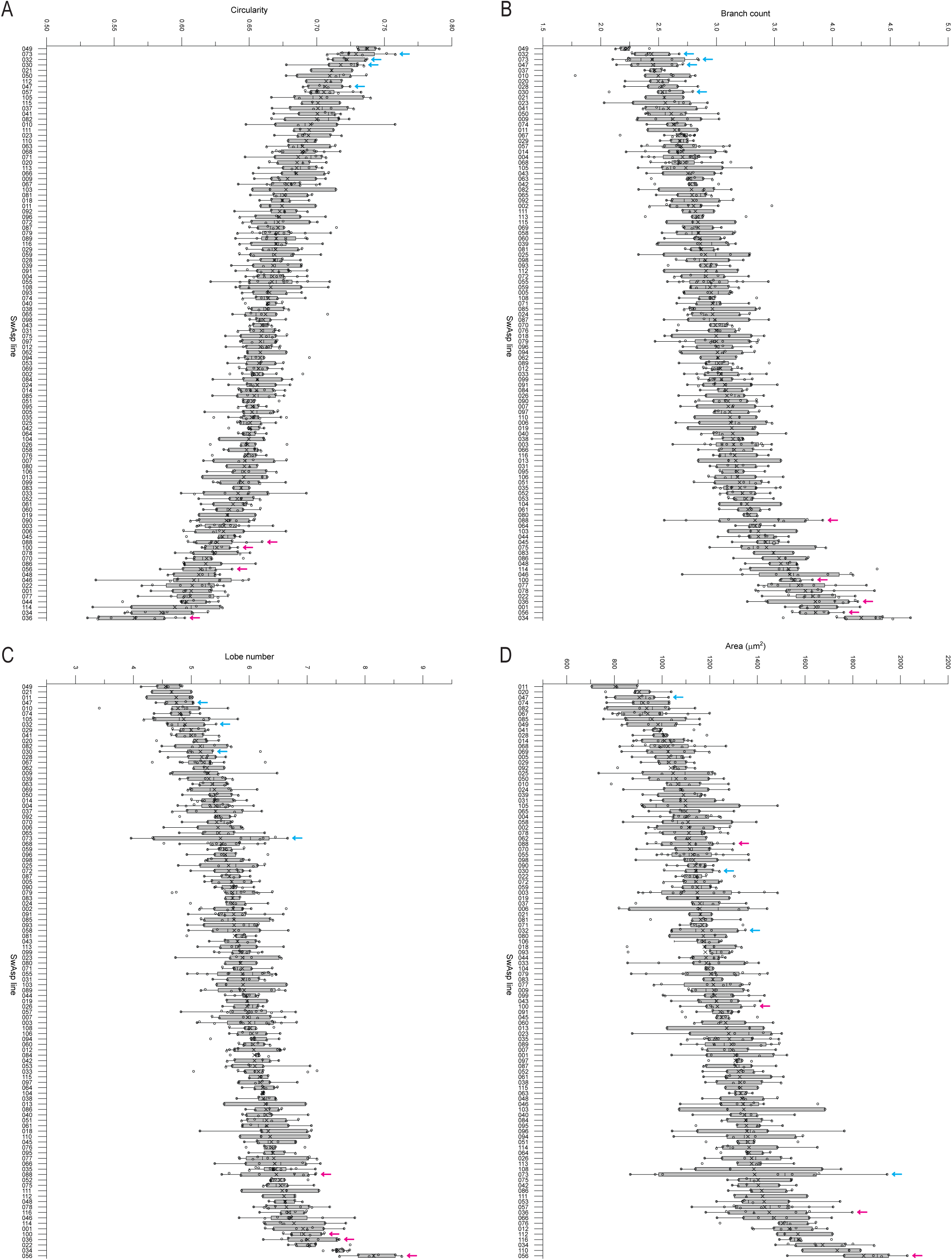
Pavement cell shape complexity varies among genotypes of Swedish aspen in the field. A-D, circularity (A), branch count (B), lobe number (C) and area (D) of pavement cells in adaxial epidermis of pre-formed leaf in adult trees from the SwAsp collection of European aspen genotypes grown in the field. To ease visualization, each data point in the box plot represents the mean value of 70-160 cells measured per leaf rather than showing each measured cell; 2-3 leaves per tree and 1-5 trees per genotype were analyzed. Those genotypes selected as representatives of more and less complex cell shape for further studies with terminal leaves of juvenile trees are indicated with magenta and cyan arrows, respectively.

Since there is a reference genome assembly for European aspen^17,18^ and the genomes of most of the SwAsp genotypes have been fully resequenced^19^, we took a GWAS approach with the aim of uncovering single nucleotide polymorphisms (SNPs) significantly associated with the cell shape traits. In this way, we identified a T-to-C SNP (chr8_11748682_C_T) in the sequence of a MYB transcription factor gene, *MYB305a* (Potra2n8c18226), that was significantly associated with increased branch count with additive effects (Fig. 2A and Fig. S3A). This SNP explained 30% of the variation in branch count across the genotypes included in the GWAS and besides being the highest-ranked SNP associated with branch count in terms of *P*-value, the candidate SNP was ranked number 233 for circularity, 685 for lobe number, 23 for solidity and 11 for lobeyness (Supplementary Data S2). Among the size-related features, the candidate SNP ranked number 120 for perimeter and 2805 for length and was not ranked among the SNPs in the top 1000 genes for non-lobe area, cell area or width (Supplementary Data S2). The majority of the SwAsp genotypes were homozygous for the reference T allele for the candidate SNP, and they were generally classified in a less complex category of pavement cell shape, with lower branch count, lobe number and lobeyness, and with higher circularity and solidity than the other genotypes (Fig. 2B and Fig. S3B-E). Another large group of genotypes were heterozygous for the T-to-C allele, and they could generally be seen to display higher branch count, lobe number and lobeyness, as well as lower circularity and solidity than the previous category (Fig. 2B and Fig. S3B-E). Finally, two genotypes were homozygous for the alternative C SNP allele, and these were among the most complex cell shape genotypes, with among the highest branch count, lobe number and lobeyness values, and among the lowest circularity and solidity values of all the genotypes (Fig. 2B and Fig. S3B-E). For the size-related features, including non-lobe area, cell area, perimeter, length and width, the genotypes tended to overlap more than for the shape-related features, although the two C/C genotypes did have somewhat larger cells than many of the other genotypes (Fig. S3F-J). These results suggest that *MYB305a* may be involved in regulating pavement cell shape in European aspen leaves.

**Fig. 2:**
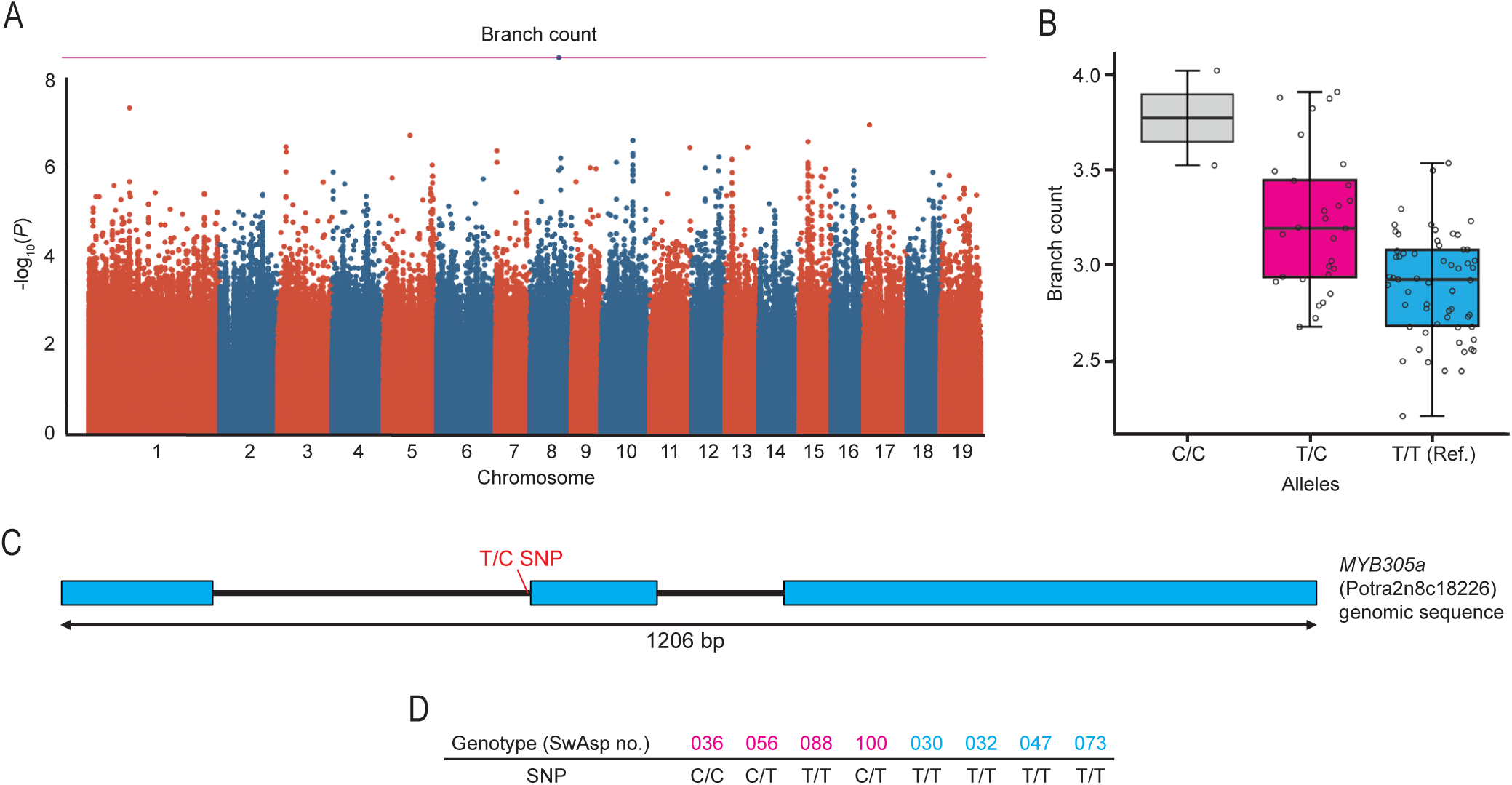
Identification of a SNP in *MYB305a* associated with pavement cell shape complexity in aspen. A, manhattan plot of the *P*-values obtained for branch count in the GWAS, in which a T-to-C SNP in the transcription factor *MYB305a* (Potra2n8c18226) was identified as being significantly associated with pavement cell branch count among the SwAsp collection of European aspen genotypes grown in the field. B, branch count of pavement cells in the SwAsp genotypes analyzed, grouped according to the alleles of the candidate SNP present in the genotypes. C, position of the candidate SNP in the first intron of the *MYB305a* gene in European aspen. Introns are shown as black lines and exons as cyan boxes. D, alleles of the candidate SNP in the SwAsp genotypes selected as representatives of more (in magenta) and less (in cyan) complex cell shape.

The broad-sense heritability of all the pavement cell traits was high, especially so for most of the shape-related features (Fig. S4A), indicating high power for the GWAS to detect associations even for small-effect alleles, and suggesting that environmental effects on the traits were relatively small. Genetic correlations of the cell features revealed the presence of two distinct axes of variation, visualized by two blocks in the heat map of genetic correlations – one representing shape simplicity traits (solidity and circularity) and the other representing shape complexity and size traits (all other shape-related features and all size-related features) (Fig. S4B). While solidity and circularity were strongly positively correlated with each other, they were somewhat negatively correlated with the size-related traits. Intuitively, circularity and solidity were strongly negatively correlated with the shape-related traits lobe number, branch count and lobeyness. In turn, lobe number, branch count and lobeyness were somewhat positively correlated with the size-related traits (Fig. S4B). Overall, there is a genetic correlation between pavement cell size and shape, with more complex cells tending to be larger.

The identified candidate SNP [chr8_11748682_C_T] is located at the end of the first intron of *MYB305a* (Fig. 2C) and the T-to-C variation is present in three of the four previously selected representative genotypes for more complex pavement cell shape, while being absent from all four representatives for less complex shape (Fig. 2D). To investigate whether the SNP may interfere with RNA splicing, we measured expression of *MYB305a* in the representative SwAsp genotypes for more and less shape complexity with two primer pairs, one of which spanned the first intron and one of which did not span an intron (Fig. S5A) and compared the results. For the primer pair that spanned the first intron, a product of the expected size of about 200 bp was amplified from cDNA in all lines, although additional, larger products were also amplified in several of the lines, possibly representing alternative transcripts (Fig. S5B). The relative gene expression levels measured by RT-qPCR using the other primer pair (Fig. S5C) generally matched the abundance of the product detected using the primer pair spanning the first intron (Fig. S5B), suggesting that the candidate SNP does not interfere with RNA splicing. Many transcription factors contain regulatory elements in their introns, as exemplified by the floral identity transcription factor gene *AGAMOUS* in Arabidopsis^20^, therefore it is plausible that the location of the candidate SNP is important for *MYB305a* regulation, although the gene expression level did not appear to correlate with the SNP allele (Fig. S5C and Fig. 2D). The closest orthologs of *MYB305a* in Arabidopsis *MYB71* (also known as *MYB305*; AT3G24310) and *MYB79* (AT4G13480) (Fig. S5D), which are nuclear-localized transcriptional activators that show enhanced expression in response to abscisic acid (ABA) treatment^21^. Furthermore, *MYB71* has been shown to be involved in ABA and abiotic stress responses^21^.

### Increased complexity of pavement cell shape from field to greenhouse conditions

To investigate the robustness of pavement cell shape complexity in the SwAsp lines grown in different conditions, leaves from juvenile trees (so-called terminal leaves formed at the terminal meristem and emerging directly from the stem) grown in controlled greenhouse conditions were used. We analyzed pavement cell shape complexity and area in terminal leaves of different developmental stages for the eight SwAsp genotypes previously selected as representatives of more and less complex pavement cell shape. Adaxial pavement cell shape complexity increased, with circularity decreasing and both branch count and lobe number increasing, over the terminal leaf series as the leaves developed and the cells grew in area (Fig. 3A-D). By leaf 17, there was a clear separation of the genotypes into two groups of more or less complex cell shape for all the shape features measured (Fig. 3A-C and Fig. S6A-H). Moreover, the lines followed the same trend in shape complexity for terminal leaves as was found for pre-formed leaves, with those genotypes that were the most or least complex for pre-formed leaf pavement cell shape also being the most or least complex for pre-formed leaf pavement cell shape, respectively (Fig. 3A-C compared to Fig. S1A-C), indicating high robustness of these traits. However, for terminal leaf 17, in which the cells were almost fully expanded for most of the genotypes, the pavement cells were generally more complex in shape, with lower circularity and higher branch count and lobe number, than in the pre-formed leaves of the same SwAsp genotypes in the field (Fig. 3A-C and Fig. S6A-H compared to Fig. S1A-C and Fig. S2A-H). Overall, we concluded that juvenile trees grown in the greenhouse were suitable for further experiments investigating the role of *MYB305a* in European aspen leaf pavement cell shape regulation. Interestingly, despite being simpler in shape, pavement cell area was larger in the pre-formed leaves from trees in the field than in mature terminal leaves from juvenile trees in the greenhouse for all genotypes except SwAsp100 (Fig. 3D compared to Fig. S1D).

**Fig. 3:**
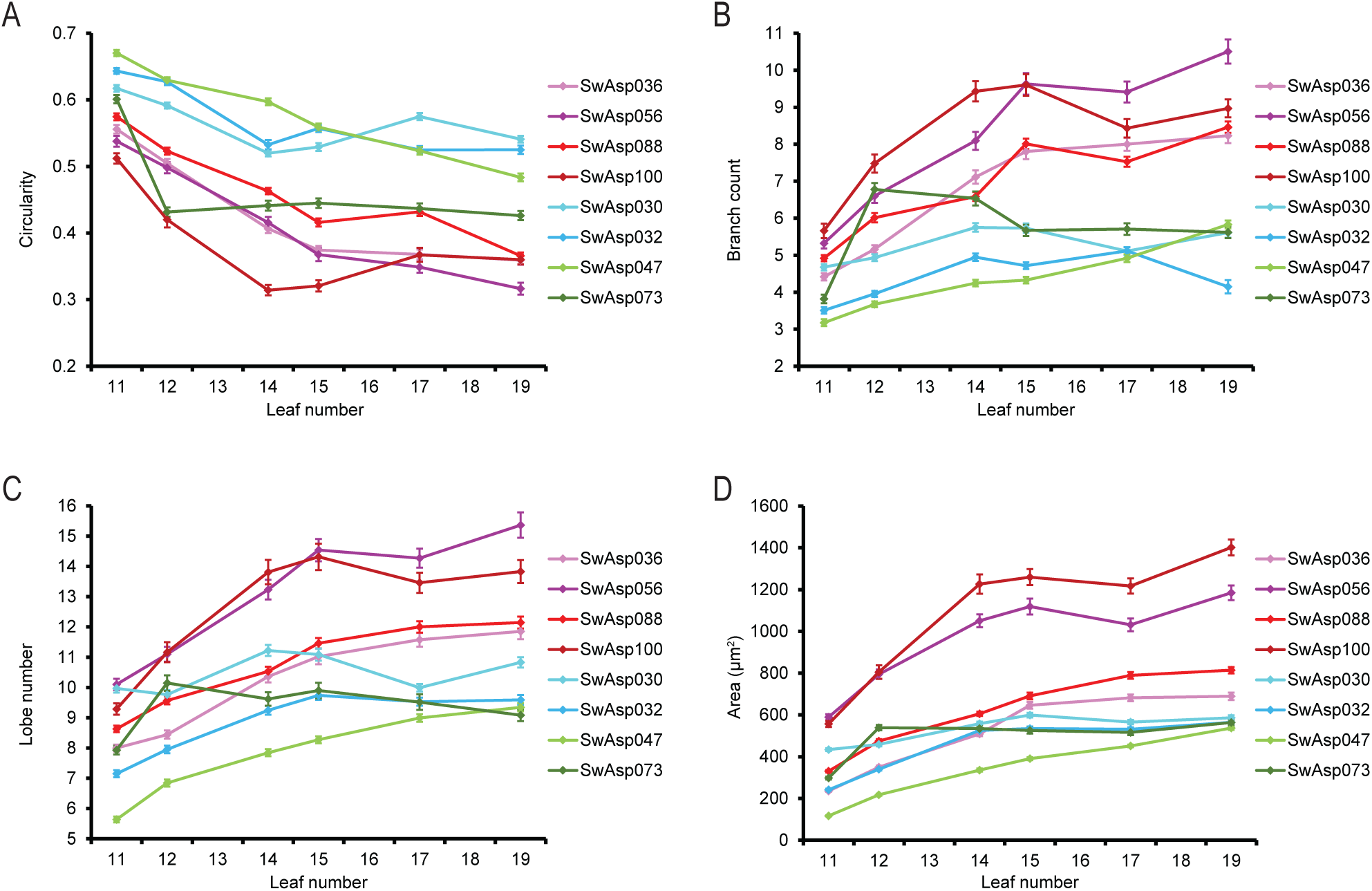
Pavement cell shape complexity in terminal leaves of juvenile aspen in the greenhouse. A-D, circularity (A), branch count (B), lobe number (C) and area (D) of pavement cells in adaxial epidermis of terminal leaf numbers 11-19 in juvenile trees from the SwAsp collection of European aspen genotypes grown in the greenhouse. Leaf number was counted from the top down, with number 1 being the youngest leaf. Each data point represents the mean value of 30-250 cells in total, measured across 1 leaf per tree and 3 trees per genotype. Error bars represent standard error of the mean. SwAsp genotypes from the more and less complex pre-formed leaf pavement cell shape groups are show in magenta/red and cyan/green colors, respectively.

### Genetic modifications of *MYB305a* suggest its involvement in pavement cell shape regulation

To validate the GWAS results and confirm a role for *MYB305a* in pavement cell shape regulation in European aspen leaves, we intended to generate *MYB305a* native promoter-driven GFP reporter lines, CRISPR/Cas9-generated *MYB305a* large-fragment deletion mutants and *MYB305a* over-expressor lines. We first attempted to generate these lines using SwAsp036 and 030 as the genetic backgrounds, representing genotypes with the most and least complex cell shapes, respectively. However, these lines were not amenable to any of these genetic manipulations since no plants survived the procedures. We therefore instead turned to clone T89 of hybrid aspen (*Populus tremula x tremuloides*), which is known to be easily transformed. T89 is homozygous for the reference T SNP allele of *MYB305a* (Potrx042905g12730) at the position of interest. We first confirmed that developing terminal leaves from juvenile T89 trees in the greenhouse display similar progression of cell shape complexity acquisition and cell growth during leaf development as the SwAsp genotypes (Fig. S7A-D compared to Fig. 3A-D). We then proceeded with our intended genetic modifications in T89 and were successful in generating all of the desired lines.

We obtained six *ProMYB305a:GFP* lines in T89 and used confocal microscopy imaging to investigate GFP fluorescence in the terminal leaves. In three of the lines, we were not able to visualize any GFP fluorescence, while in two of the lines, we observed very faint GFP fluorescence in the adaxial pavement cells (Fig. S8A). However, in one line, number 4, we found stronger, more clearly visible GFP fluorescence in the pavement cells (Fig. S8A). Together, the imaging of the various lines suggested that while the *MYB305a* promoter is indeed expressed in adaxial pavement cells of terminal aspen leaves, it is likely expressed at a rather low level. We next investigated the GFP expression patterns in a series of terminal leaves of different developmental stages in juvenile trees of *ProMYB305a:GFP* line 4 grown in the greenhouse. While the expression level was very low in younger leaves, numbers 9-13, it was not consistent throughout the epidermis, with clearly enhanced expression being observed in specific pavement cells across the tissue (Fig. 4A-C). A striking increase in expression was observed across the epidermis in the older leaves, numbers 15-19 (Fig. 4D-F), compared to the younger leaves.

**Fig. 4:**
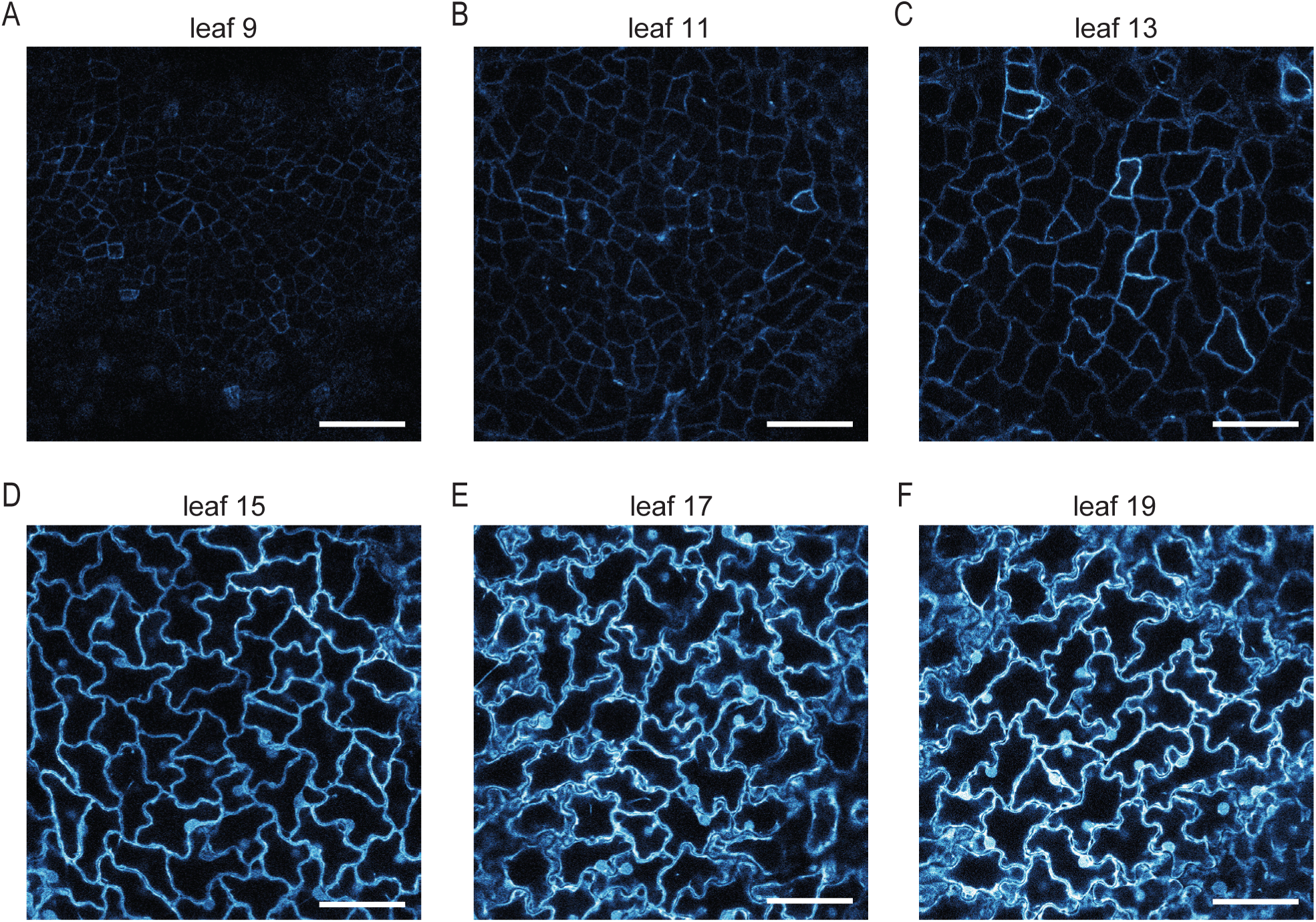
*MYB305a* promoter expression increases in pavement cells as they grow and acquire their complex shape. A-F, representative confocal microscopy images of GFP fluorescence in pavement cells in adaxial epidermis of terminal leaf numbers 9-19 in *ProMYB305a:GFP* line 4 of hybrid aspen (T89) grown in the greenhouse. Increasing GFP fluorescence is represented by a cyan-to-white color scale. Scale bars represent 50 μm.

Using CRISPR/Cas9 technology, we generated *MYB305a* large-fragment deletion lines in T89 and identified six lines in which deletions of 371-396 bp in both alleles were confirmed by sequencing (Fig. S8B-C). We also generated five over-expressor lines of *MYB305a* in T89 and confirmed highly variable *MYB305a* over-expression levels in these lines by RT-qPCR in terminal leaves (Fig. S8D). During the RT-qPCR experiments, we noticed that *MYB305a* expression was generally rather low in the leaf blade tissue of T89 controls, with amplification becoming detectable only after 30 amplification cycles (compared to 19 cycles for the house-keeping gene). For the four deletion lines that we grew in the greenhouse, *myb305a* line 4, 20, 24 and 29, we did not observe any significant difference in shoot height compared to T89 after 8 weeks of growth in soil (Fig. S9A-B). Of the over-expressor lines, *Pro35S:MYB305a* line 1, which had the highest over-expression level of around 54,000-fold (Fig. S8D), was severely stunted in shoot growth after 8 weeks of growth in soil, with much smaller leaves and significantly shorter shoots than T89 (Fig. S9C-D). We therefore decided not to work further with this line. None of the other *Pro35S:MYB305a* lines were affected in shoot height, but line 5, which displayed the next-highest over-expression level of around 11,000-fold (Fig. S8D), had somewhat smaller leaves than T89 (Fig. S9C-D).

We then selected three of the deletion lines, *myb305a* line 4, 20 and 24, and three of the over-expressor lines, *Pro35S:MYB305a* line 3, 4 and 5, for studies of adaxial pavement cell shape in terminal leaves of juvenile plants grown in the greenhouse. Compared to T89, *myb305a* line 4 and 20 displayed slightly but significantly increased circularity, decreased branch count and decreased lobe number in the pavement cells, while line 29 was unaffected in these features compared to T89 (Fig. 5A-C). Interestingly, *myb305a* lines 20 and 29 had deletion lengths expected to result in frameshift mutations, while line 4 did not, which suggests not only that MYB305a plays a role in regulating pavement cell shape in aspen, but also that the deleted region itself is important for this function of MYB305a. The three *Pro35S:MYB305a* lines all showed significant increases in circularity and decreases in branch count and lobe number compared to T89 and these effects were generally somewhat stronger than those observed in the *myb305a* lines (Fig. 5A-C). Finally, the decreased cell shape complexity in these lines was generally not related to any change in cell size, since cell area was only significantly affected in one of the *myb305a* lines, in which it was slightly decreased (Fig. 5D). Together, these results strongly suggest that *MYB305a* is involved in pavement cell shape regulation in aspen leaves. Since both *myb305a* and *Pro35S:MYB305a* lines generally displayed decreased pavement cell shape complexity without cell area being affected, we could conclude that precise, fine-tuned levels of MYB305a must be important for promoting cell shape acquisition independently from general cell growth regulation.

**Fig. 5:**
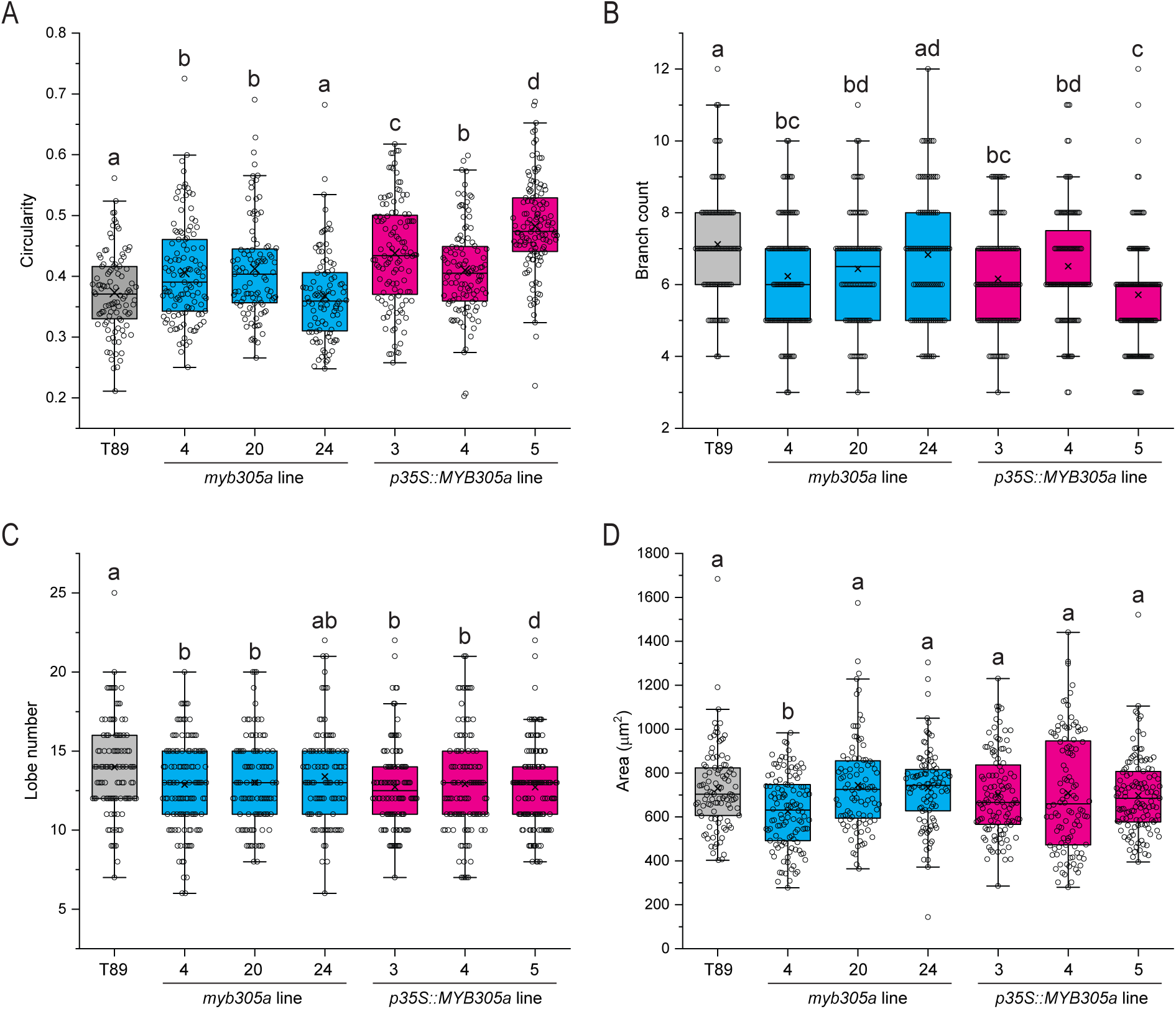
Pavement cell shape features of *MYB305a* CRISPR mutant and over-expressor lines. A-D, circularity (A), branch count (B), lobe number (C) and area (D) of pavement cells in adaxial epidermis of terminal leaf number 17 in *MYB305a* CRISPR mutant (*myb305a*, in cyan) and over-expressor (*Pro35S:MYB305a*, in magenta) lines of hybrid aspen (T89) grown in the greenhouse. Each data point in the box plot represents one of 95-125 cells in total measured across 1 leaf per tree and 3 trees per genotype. Different letters indicate significantly different distributions at *P* < 0.05 (Wilcoxon rank sum test).

There is a close homolog of *MYB305a* in European aspen, Potra2n10c21884, which we called *MYB305b* (Fig. S5D) and which corresponds to Potrx058201g19640 in hybrid aspen. We therefore also attempted to generate double *MYB305a MYB305b* CRISPR/Cas9 large-fragment deletion lines in T89. Using the same strategy as before, with the same guide RNAs, we were successful in generating large-fragment deletions or inversions in *MYB305a* in most of the lines we generated, as confirmed by sequencing (Fig. S10A). However, our attempts to generate large-fragment deletions in *MYB305b* (Fig. S10B) were generally unsuccessful (Fig. S10C). We did, however, confirm one-nucleotide indels around the sites of expected DNA breaks in most of the lines (Fig. S10C). For the five lines that we grew in the greenhouse, *myb305ab* line 25, 26, 29, 32 and 36, we observed an apparent slight decrease in shoot height after 8 weeks of growth in soil, but there were no statistically significant differences to T89 shoot height (Fig. S10D-E). We randomly selected three of the lines, *myb305ab* lines 26, 32 and 36, in which we confirmed a large-fragment deletion or inversion in both alleles of *MYB305a* and expected a frameshift mutation in both alleles of *MYB305b*, for studies of adaxial pavement cell shape in terminal leaves of juvenile plants grown in the greenhouse. Compared to T89, two of the *myb305ab* lines displayed slightly but significantly increased circularity in the pavement cells, with line 36 being the most affected (Fig. S11A). The same two lines displayed significantly decreased branch count and line 36 additionally displayed significantly decreased lobe number and cell area (Fig. S11B-D). These results were similar to those found for the *myb305a* lines and the lack of an additive effect in *myb305ab* compared to *myb305a* suggests that *MYB305b* might not play a functionally redundant role with *MYB305a* in the regulation of pavement cell shape in aspen leaves.

### *MYB305a* is drought stress-responsive and may regulate pavement cell shape response to abiotic stress

The closest homolog of aspen *MYB305a* in Arabidopsis, *MYB71* (also known as *MYB305*; AT3G24310) (Fig. S5D), has been shown to be up-regulated in response to abscisic acid treatment and to positively regulate a number of stress response-related genes^21^. Since abscisic acid is known to mediate drought stress responses in plants^22^ and the expression of many other MYB transcription factors is regulated by drought^14^, we planned to test whether drought stress would affect *MYB305a* expression in T89 terminal leaves. After a moderate drought stress treatment of several days on *ProMYB305a:GFP* line 4, which resulted in a significant reduction in shoot height and some leaf wilting in the lower half of the shoot (Fig. S12A-B), the epidermis in terminal leaves of different ages was imaged with confocal microscopy in the drought-stressed trees as well as non-drought controls. In leaf numbers 9-13, in which the cells developed from rather small and square-shaped to much larger with more complex shape, a clear enhancement of GFP fluorescence was visible in the drought-stressed trees compared to the control trees (Fig. 6A-C). Simultaneously, the drought stress also induced a significant increase in both cell area and shape complexity compared to the controls in these younger leaves (Fig. S12C-F). Surprisingly, once GFP fluorescence had reached around the same levels in the controls as in the drought treatments, from around leaf 15 and upwards (Fig. 6D-F), the drought stress began to have the opposite effect on cell size and shape, resulting in a significant reduction in cell area and shape complexity compared to the controls (Fig. S12C-F). These results imply that *MYB305a* promoter expression is strongly induced by drought stress in younger aspen leaves and that *MYB305a* may be involved in enhancing the growth and development of the pavement cells in these young leaves in response to the drought stress. However, in older leaves, in which *MYB305a* promoter expression is already much higher than younger leaves in the controls, the later stages of pavement cell growth and development are repressed by the drought stress. These results agree with the idea that fine-tuned MYB305a levels are important for regulating the correct pavement cell shape complexity level according to the stage of development and environmental conditions.

**Fig. 6:**
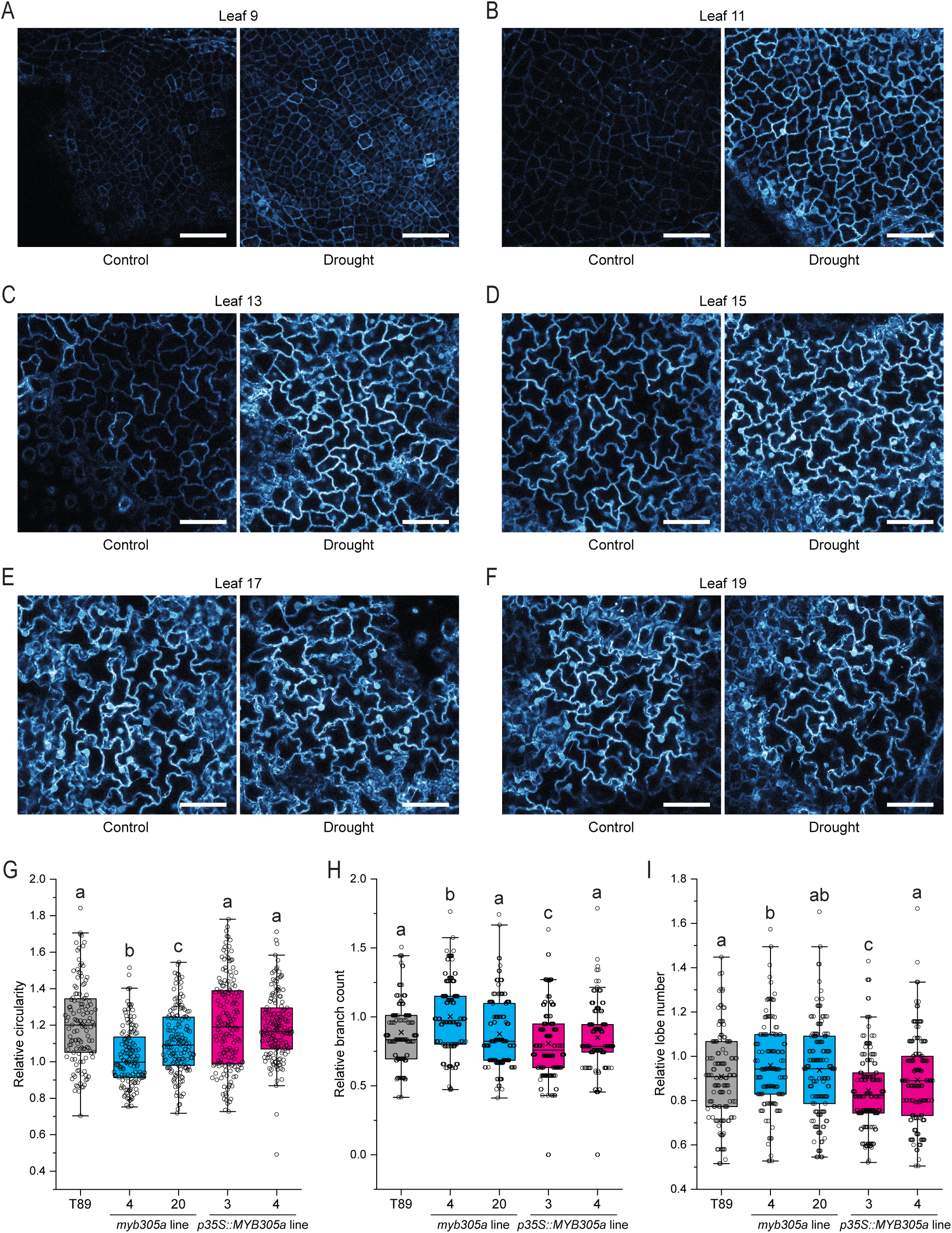
*MYB305a* may play a role in the response of pavement cell shape to drought stress. A-F, representative confocal images of GFP fluorescence in pavement cells in adaxial epidermis of terminal leaf numbers 9-19 in *ProMYB305a:GFP* line 4 of hybrid aspen (T89) grown in control or drought conditions in the greenhouse. Increasing GFP fluorescence is represented by a cyan- to-white color scale. Identical acquisition settings were used for each image. Scale bars represent 50 μm. G-I, relative drought:control circularity (G), branch count (H) and lobe number (I) of pavement cells in adaxial epidermis of terminal leaf number 17 in *MYB305a* CRISPR mutant (*myb305a*, in cyan) and over-expressor (*Pro35S:MYB305a*, in magenta) lines of hybrid aspen (T89) grown in the greenhouse. Relative cell shape features were calculated as the drought:control ratio; each data point in the box plot represents the ratio of one of 125-160 cells in total measured across 1 leaf per tree and 3 trees per genotype in drought stress conditions, compared to the mean control (no drought stress) value. Different letters indicate significantly different distributions at *P* < 0.05 (Wilcoxon rank sum test).

To investigate the idea that *MYB305a* may be involved in the responses of pavement cell shape to the drought stress, we next tested the effects of the drought treatment on cell shape features in two of each of our *myb305a* deletion and *Pro35S:MYB305a* over-expressor lines. The drought treatment resulted in significantly reduced shoot height in all the lines as well as in T89 control trees (Fig. S13A-B). We compared any drought stress-induced changes in cell shape features in leaf 17 of the transformed lines to that of T89. In T89, pavement cell circularity was increased 1.2-fold by the drought stress compared to control conditions (Fig. 6G). Interestingly, both of the *myb305a* lines were resistant to this effect of drought stress on circularity, while the *Pro35S:MYB305a* lines were affected in circularity to the same degree as T89 (Fig. 6G). In T89, the drought stress resulted in around 0.9-fold decreases in pavement cell branch count and lobe number compared to control conditions (Fig. 6H and I). While one of the *myb305a* lines was affected to a similar extent as T89, the other *myb305a* line was resistant to this effect of drought stress on both branch count and lobe number (Fig. 6H and I). Conversely, while one of the *Pro35S:MYB305a* lines was affected to a similar extent as T89, the other *Pro35S:MYB305a* line was over-sensitive to the effect of drought stress on branch count and lobe number, in that they were reduced more than in T89 (Fig. 6H and I). Although these results are not unequivocal, taken together with the effects of drought stress on inducing *MYB305a* promoter expression, they imply that *MYB305a* may play a role in the response of pavement cell shape complexity to drought stress in aspen leaves.

We next turned our attention back to the SwAsp population, since we showed that the genotypes naturally vary in pavement cell shape complexity and earlier work has estimated the water-use efficiency of the different genotypes^23^. We performed *k*-means clustering to partition the data into an optimal number of clusters and observed that our selected representative genotypes of more and less complex pavement cell shape were divided between the two resulting clusters (Fig. S14A). To investigate whether the two clusters really do divide the genotypes into groups of more and less complex pavement cell shape, we color-coded the branch count values of the genotypes measured earlier according to their clusters (Fig. S14B). This indicated that the genotypes were generally separated according to pavement cell shape complexity by the clustering. We therefore statistically compared the water-use efficiency of the genotypes (estimated by a measure of the foliar carbon isotope ratio or delta^13^C) between the two clusters (Fig. S15A). Although the difference between the clusters was not statistically significant, the rather low *P*-value suggests the possibility of a mild negative correlation between pavement cell shape complexity and water-use efficiency (Fig. S15A). Records of environmental variables at the original SwAsp sampling sites are available, including precipitation in the driest month of the year and annual precipitation, both of which we averaged over a 30-year period ending just 3 years before the genotypes were originally sampled. We compared the average precipitation in the driest month of the year between the clusters and once again found a non-significant difference that nevertheless had quite a low *P*-value (Fig. S15B). These results suggest a mild link between pavement shape complexity and precipitation, with the more complex cell shape cluster genotypes tending to originate from sampling sites with less precipitation in the driest month (Fig. S15B). We next compared the average annual precipitation between the clusters and this time we found a highly significant difference, with the more complex cell shape cluster genotypes tending to originate from sites with less precipitation (Fig. S15C). Finally, we compared both the latitude and the longitude of the original sampling site between the clusters and found highly significant differences for both, with the more complex cell shape cluster genotypes tending to originate from more northern and more eastern sampling sites (Fig. S15D-E). These results strengthen the idea that climatic variables affect the shape complexity of leaf pavement cells in the genotypes. Taken together, our work suggests that the level of pavement cell shape complexity in aspen leaves results from a combination of evolutionary adaptation to environmental conditions such as water availability within a genotype, and developmental responses to local stresses such as drought in the individual tree during leaf growth. Furthermore, our results provide a first step towards unravelling the molecular mechanisms controlling leaf pavement cell shape in aspen in response to environmental conditions, by implicating the involvement of the transcription factor MY305a.

## Discussion

Although the mechanisms for regulation of pavement cell shape complexity in aspen leaves are largely unknown, our work has provided an important first step in understanding this process by revealing the involvement of the transcription factor MYB305a. This was facilitated by access to the SwAsp collection of Swedish aspen genotypes, which allowed us to link a biological process to a natural population originating from native habitats, rather than an artificial environment. We also detected correlations between the locations and environments of the sampling sites of the SwAsp clones and the complexity of pavement cell shape. In particular, our results as a whole suggest a strong link between water use/availability and pavement cell shape. At first glance, the negative correlation of water use efficiency and precipitation with cell shape complexity may seem to contradict with the effect of drought treatment on simplifying cell shape. However, the SwAsp experiments were performed with pre-formed leaves sampled from adult trees in the field, while the drought experiments were performed with terminal leaves from greenhouse-grown juvenile trees, with the different environments and leaf types likely to affect the results. This apparent contradiction was also mirrored in the drought experiment on T89 leaves of different developmental stages, which surprisingly showed drought-induced enhancement of shape complexity in younger leaves but simplification of cell shape complexity in older leaves. While these results suggest that pavement cell shape responds to water stress, there are likely to be many additional factors influencing the final cell shape.

We found strong differences in the level of pavement cell shape complexity between pre-formed and terminal leaves. These two leaf types from the same species differ considerably in size and shape and although they are often associated with adult and juvenile trees, respectively, they can also grow on the same shoot^24^. Such heterophylly is common among *Populus* species and may be linked to adaptations to environment or growth conditions, as exemplified by the desert riparian poplar (*P. euphrates*). This species forms narrow leaves at the bottom of the canopy, with the leaves becoming more ovate the higher up in the canopy they grow, which is thought to be an adaptive feature and has been linked to water availability and salt stress response^25–27^. Interestingly, similarly to our findings for field-grown and greenhouse-grown European aspen, pavement cells in camphor tree (*Camphora officinarum*) were recently also shown to display much more complex, lobed shapes when the plants were grown indoors compared to when grown outdoors^3^. Additionally, many plant species have been found to display widely different pavement cell morphologies in different growth conditions and/or different organs of the same plant^3,6,13^, suggesting that environmental factors are likely to influence pavement cell shape development. In European aspen, fully developed terminal leaves are considerably larger in area than fully developed pre-formed leaves^24^ and larger leaves may be expected to result in more complex pavement cell shapes as a strategy to mitigate the mechanical stresses imposed by the more extensive near-isotropic expansion in larger organs^3^.

Over the last few years, the SwAsp collection has proven itself to be a highly valuable resource for tree-related research in Sweden, convenient for investigating a diverse range of biological questions^15,28–32^. The GWAS approach itself has proven a powerful strategy to reveal new molecular players in regulatory processes in *Populus* species. For example, using GWAS for measures of stem growth traits over time in a large population of Chinese white poplar (*P. tomentosa*) genotypes, a role for prolyl 4-hydroxylase 9 was revealed and subsequently confirmed in the promotion of stem diameter growth^33^. A GWAS strategy was also successfully employed in natural populations of two *Populus* species, Chinese white poplar and Chinese cottonwood (*P. simonii*), to reveal new molecular players in *Populus* leaf morphology^34^. On the other hand, GWAS for leaf shape features in the SwAsp genotypes of European aspen failed to reveal statistically significant SNP associations, which combined with further gene expression analyses suggested that leaf shape variation is a complex, omnigenic trait in aspen^24^. Similarly, no SNPs across the SwAsp genotypes were found to be significantly associated with total fatty acid content of stems, suggesting that this trait is highly polygenic^35^. In our study, we only found a single SNP that was significantly associated with pavement cell shape in the SwAsp genotypes, and only for one cell shape trait across all that we included, suggesting that leaf pavement cell shape in European aspen may also be highly polygenic. While quantitative trait loci (QTLs) for epidermal cell area in leaves of black cottonwood (*P. trichocarpa*) and eastern cottonwood (*P. deltoides*) have been identified previously^36^, we are not aware of any work identifying QTLs or significant SNP associations by GWAS for shape features in *Populus* epidermal leaf cells. Importantly, we succeeded in subsequent validation of our single significant SNP using transgenic approaches, supporting a role for *MYB305a* in aspen pavement cell shape regulation.

MYB305a is a member of the R2R3-type MYB transcription factors, a large family of transcriptional regulators, including both activators and repressors of gene expression, that are involved in many physiological processes^37,38^. In black cottonwood, there are almost 200 R2R3-type *MYB* genes, suggesting a diverse range of functions and implying a role for this family in the phenotypic diversity of *Populus* species^39^. Transcription factors are themselves also subject to complex expressional regulation mechanisms and contain regulatory elements in their genes, which can occur in the introns. The cell shape-associated SNP we identified through our GWAS is located in the first intron of *MYB305a*, presumably at a site particularly important for its regulation. The presence of the T-to-C mutation at the SNP site resulted in enhanced pavement cell shape complexity in the aspen genotypes, but whether this sequence change results in enhanced or reduced *MYB305a* expression is difficult to conclude, since we found that both deletion and over-expression of the gene resulted in reduced pavement cell shape complexity. The cell shape regulatory pathways in which MYB305a is active are likely to be complex, in which precise levels of this transcription factor are important for achieving the proper level of cell shape complexity needed according to the developmental context and environmental conditions. To our knowledge, *MYB305a* has not previously been studied in *Populus* species, but investigations on its close homolog in Arabidopsis, *MYB71*, revealed its function as a transcriptional activator in the nucleus, its up-regulation in response to abscisic acid treatment and its likely activity as a positive regulator of abscisic acid response^21^. Since abscisic acid is a well-known mediator of stress-responsive signaling^22,40,41^, the authors then investigated *MYB71*-induced gene up-regulation, revealing a small set of genes of which the majority are known to be involved in stress responses^21^. Many R2R3-type MYB family members are up- or down-regulated by drought treatment and thought to play roles in drought stress responses in various different species^42–49^. The up-regulated expression of several R2R3-type MYB-encoding genes in response to drought stress has recently been shown in black cottonwood and we demonstrated that *MYB305a* expression is also clearly up-regulated in European aspen leaves upon drought stress. Overall, our results suggest that this transcription factor regulates drought stress-induced changes in pavement cell shape complexity in aspen leaf epidermis.

Pavement cell shape is hugely variable both among and within plant species^6,13^ and the biological reasons for the interdigitated, jigsaw puzzle-like shape are therefore not entirely obvious. Enhanced integrity and reduced mechanical stress in the epidermis of a near-isotropically expanding organ potentially exposed to a wide variety of environmental stresses are likely functions of the jigsaw puzzle shape^1,50^ and these roles may partially explain apparent cell shape plasticity under different conditions including abiotic stress^6^. Interestingly, a set of closely related R2R3-type MYB family genes have been reported to regulate epidermal cell shape in various organs across several species. Early work suggested that *MIXTA* in snapdragon (*Antirrhinum majus*) regulates the cone shape of epidermal cells in flower petals^51^, which was later also shown for *MIXTA*-related *MYB* genes in the same and other species, including

Arabidopsis *MYB16*^52^. *MIXTA*, *MYB16* and the close homolog *MYB106* in Arabidopsis have additionally been implicated in the regulation of leaf trichome shape^53–55^. Roles for *MYB16* and *MYB106* in cuticle biosynthesis have also been shown^56^ and more recently, *MYB16* was additionally suggested to regulate the division and differentiation of young stomatal lineage-derived pavement cells in the cotyledon epidermis^57^. These studies reveal that the *MIXTA*-like R2R3-type MYB transcription factors play multi-functional roles in plant development but seem to be particularly important for regulation of epidermal cell shape. Our work specifies that *MYB305a* regulates pavement cell shape in aspen leaves, but whether this role is specific to this cell type and species is still unknown. Many questions also arise regarding which molecular signaling pathways and mechanisms *MYB305a* acts in to regulate this process and we look forward to the identification of interactors and targets of this transcription factor in aspen leaves in the future.

## Methods

### Plant material and growth conditions

For phenotyping of pavement cells from adult trees in the field, mature pre-formed leaves were harvested from genotypes of the SwAsp collection of European aspen (*Populus tremula*) maintained by Skogforsk in the common garden at Sävar, Northern Sweden, during the summer months. Only mature, undamaged, non-shaded leaves were selected for harvesting from a height of about 1.5 m above the ground, and the first and last leaves from leaf cohorts originating from the same bud were avoided. Two to three leaves from each of one to five trees, from each of 106 different genotypes, were collected in plastic bags and stored in a cool box in the field before being transported to the lab and stored at 4°C for up to several days.

For all further experiments, trees of SwAsp genotypes and hybrid aspen (*Populus tremula* x *tremuloides*) clone T89 were obtained from the *in vitro* collections of *Populus* species maintained via shoot cutting propagation by the Umeå Plant Science Centre (UPSC) Poplar Transgenics Facility. The *in vitro* plants were maintained in sterile conditions in half-strength Murashige and Skoog (MS) medium with 3.5 g/L gelrite at pH 5.6 and incubated in 22°C under an 18/6 h light/dark regime. White paper was used as shade for the first 2 weeks of growth of new cuttings. All transgenic lines were created in the T89 background and also maintained *in vitro*. For growth in soil, three-week-old rooted *in vitro* plants were potted and immediately covered with plastic bags, which were removed 2 weeks later. For experiments with SwAsp genotypes or transgenic lines, the plants were then grown in a controlled greenhouse, or in an automated conveyor network (WIWAM Conveyor, SMO, Eeklo, Belgium) in a controlled growth room at the UPSC Tree Phenotyping Platform, respectively. The trees were grown under an 18/6 h 22/18°C day/night regime with 200 µmol/m^2^/s light intensity and 60% relative humidity. In the platform, both white (FL300 Sunlight v1.1) and red (custom-made FL100 725-735 nm) LED lamps (Senmatic A/S, Søndersø, Denmark) were used and all side shoots were removed from the trees 3 and 6 weeks after potting. Excluding the drought experiments, the trees were fertilized with Rika S (SW Horto AB, Hammenhög, Sweden) every week or every 2 days, and leaves were harvested for experiments 6 or 8 weeks after potting, respectively. The trees at the platform were photographed and measured for shoot height 8 weeks after potting. For the drought experiments, trees were grown in the platform as usual for 5 weeks, during which time the pots were weighed and watered daily to a level of 1.7 kg water per kg soil (soil target humidity (TH) 1.7) in the pots. Drought conditions were then initiated by watering less each day for 10 days, reducing the TH by 0.1 daily, and reaching a final level of TH 0.7, at which point leaves were harvested. At the same time the drought was initiated, the control, non-drought trees were watered more, increasing the TH by 0.1 daily, to reach TH 1.9 after 2 days and then maintained at TH 1.9 until leaf harvesting. Upon harvesting, leaves from each of 3 trees per genotype and growth treatment were stored in paper bags at room temperature for up to several hours, except for *ProMYB305a:GFP* leaves, for which the cut petioles were inserted into solidified half-strength MS medium in jars, which were closed and stored at room temperature for up to several hours.

### Microscopy imaging of pavement cells

All microscopic observations were made on one sample per leaf, cut from the middle of the blade on a random side of the mid-vein. Epidermal cells on the adaxial leaf side were imaged at three random positions per leaf sample, avoiding those cells overlying the veins. For analysis of pavement cell shape in pre-formed leaves, the leaf samples were cut within a few days of harvesting, then fixed by incubation in 4% paraformaldehyde in 25 mM phosphate buffered saline (PBS; 8 g/L NaCl, 0.2 g/L KCl, 1.78 g/L Na_2_HPO_4_·2H_2_O, 0.24 g/L KH_2_PO_4_, pH 7.4) under a vacuum for 1 h, followed by storage overnight at 4°C. The samples were then washed and dehydrated according to the following protocol: 25 mM PBS for 10 min 3 times, 30% ethanol for 10 min 3 times, 50% ethanol for 10 min 3 times, and then stored in 50% ethanol at 4°C. Prior to microscopy imaging, the leaf samples were rehydrated by incubating for 1 h in each of the following ethanol series: 40%, 30%, 20%, 10%, before transferring to PBS. The samples were subsequently stained in 0.1% calcofluor white for 30 min, followed by three rinses with water, before transferring to PBS overnight. Samples were mounted in PBS and Z-stack imaging encompassing all visible calcofluor white fluorescence, with 2 µm distance between each slice, was performed on a Zeiss LSM 780 confocal laser scanning microscope. The ImageJ (Fiji) macro SurfCut ^58^ was used to extract a layer of signal relative to the sample surface in each Z-stack, followed by application of the ImageJ Gaussian blur tool to reduce the background noise. In those images with sufficient quality, the ImageJ plugin PaCeQuant^16^ from the toolbox MiToBo^59^ was then applied for automated segmentation of the pavement cell outlines and analysis of circularity, branch count, lobe number, solidity, non-lobe area, total area, perimeter, length, width and convex hull perimeter. Lobeyness was then calculated as the convex hull perimeter divided by the cell perimeter^1^. In those images without sufficient quality for automated segmentation, due to fungal/dirt contamination or uneven staining, the cell outlines were drawn by hand in Photoshop (Adobe) from maximum projections of the Z-stacks, before extraction and measurements of cell shape features and area with PaCeQuant.

For analysis of pavement cell shape in terminal leaves (except for *ProMYB305a:GFP* leaves), the leaf samples were cut on the day of leaf harvesting and stored in 70% ethanol at 4°C. The samples were rehydrated and cleared immediately prior to imaging, according to the following protocol: 50% ethanol for 15 min, 30% ethanol for 15 min, 10% ethanol for 15 min, PBS for 15 min, 5% NaOCl overnight, PBS for 15 min, fresh PBS. Imaging was performed on a Leica DMi8 epifluorescence microscope equipped with a camera. For imaging *ProMYB305a:GFP* leaves, each leaf sample was cut from the leaves stored in jars immediately prior to mounting in PBS and GFP fluorescence was imaged on a Zeiss LSM 880 confocal laser scanning microscope. For all terminal leaf microscopy images, the cell outlines were drawn by hand, extracted and the cell shape features and area measured as described above.

### Genome wide association study (GWAS)

Median values per leaf sample were calculated for each cell shape and size feature - circularity, branch count, lobe number, solidity, lobeyness, non-lobe area, cell area, perimeter, length and width to obtain one value per phenotype and biological replicate. Data for each phenotype were then parsed through a pipeline for data quality control and the calculation of a best linear unbiased predictor (BLUP) of the random effects that was subsequently used in the GWAS, as described previously^18^. The GWAS and downstream analyses of the results were performed as detailed previously^18^. Briefly, for each of the shape and size features, phenotypic BLUP values for each European aspen genotype and 6,806,717 single nucleotide polymorphisms (SNPs) were analyzed using linear mixed models in GEMMA 0.98.1^60^ with two covariates: a genetic relatedness matrix, and the latitude of origin of the SwAsp genotype to avoid confounding growth effects^18,19^. SNPs with minor allele frequency (MAF) < 0.05, Hardy-Weinberg equilibrium *P-*value < 10e-6, and missingness > 0.1 were removed prior to the analyses. The GWAS was conducted using 92 unrelated European aspen genotypes for which both phenotypic data and SNP data were available. Data were prepared and plotted in the R statistical computing environment^61^, using the qqman package for manhattan and QQ plots^62^ and the ggplot2 package for allele boxplots^63^. Broad-sense heritability was estimated using the ‘heritability’ package in R^64^. Genetic correlations were estimated as described previously^18,24^.

### Vector construction

See Table S1 for all primer sequences. To construct *ProMYB305a:GFP*, a 2.14 kb promoter region upstream of *MYB305a* was cloned from the European aspen SwAsp036 genotype using the primers proPtMYB305a_F_attB1/R_attB2 and ligated into pFASTG04 to drive GFP-GUS expression. To construct *Pro35S:MYB305a*, the 1.204 kb coding sequence of *MYB305a* without the stop codon was cloned from the European aspen SwAsp030 genotype using the primers PtMYB305a_CDS_F_attB1/R_attB2 and ligated into pFASTR05 to fuse with GFP, driven by the 35S promoter. A CRISPR/Cas9 approach based on Xing et al.^65^ was used to generate the *myb305a* single and *myb305ab* double mutants. sgRNA cassettes targeting *MYB305a* and/or *MYB305a*/*b* were amplified from pCBC-DT1T2 and/or pCBC-DT1T2/DT2T3/DT3T4 using the primer pairs PtMYB305a_sgRNA1_T1F/2_T2R and/or PtMYB305a_sgRNA1_T1F/DT1-R, PtMYB305a_sgRNA2_T2F/DT2-R, and PtMYB305b_sgRNA1_T3F/3_T4R, and ligated into pHSE401 using the Golden Gate reaction based on the following procedures: 2 μl of pHSE401, 2-6 μl of sgRNA cassette fragment, 1.5 μl of 10x FastDigest buffer (Thermo Fisher Scientific), 1.5 μl of 10 mM ATP, 1 μl of 30 U/μl T4 DNA ligase (Thermo Fisher Scientific), and 1 μl of FastDigest Eco31I (Thermo Fisher Scientific), incubated at 37°C for 5 h, then 50°C for 5 min and finally 80°C for 10 min. The vectors were cloned into chemically competent TOP10 *Escherichia coli* and purified from transformants selected on 50 µg/ml spectinomycin (for pFAST vectors) or kanamycin (for pHSE401) using the QIAprep Spin Miniprep Kit (Qiagen), according to manufacturer’s instructions. The amplified plasmids were then sent to Eurofins Genomics for sequencing.

### Plant transformation

The transformation constructs were introduced into *Agrobacterium tumefaciens* strain GV3101 by electroporation, followed by growth in liquid culture with lysogeny broth (LB) medium at 28°C with constant rotation. Antibiotic concentrations used for *Agrobacterium* selection were 20 µg/ml rifampicin, 30 µg/ml gentamicin, 10 µg/ml tetracyclin and the relevant bacterial resistance antibiotic (100 µg/ml spectinomycin or 50 µg/ml kanamycin). *Agrobacterium* pellets were obtained once the culture OD at 600 nm reached 0.3-0.8 and transformation was then performed on T89 by the UPSC Poplar Transgenics Facility. The pellets were resuspended in full-strength MS medium at pH 5.8 supplemented with 20 µM acetosyringone, to which 100-200 sterile stem and petiole pieces of 5-10 mm in length were added in petri dishes and incubated for 1 hour. All subsequent steps were performed in sterile conditions. The plant pieces were transferred to plates of MS1 medium (full-strength MS medium with 2% sucrose, 0.07% 4-morpholineethanesulfonic acid (MES), 0.2 µg/ml 6-benzylaminopurine (BAP), 0.1 µg/ml Indole-3-butyric acid (IBA), 0.01 µg/ml thidiazuron (TDZ) and 0.6% plant agar) at pH 5.8 and incubated in darkness for 2 days at 22°C. The explants were then washed twice with 500 µg/ml cefotaxime before transferring to plates of MS1 medium at pH 5.6 supplemented with 500 µg/ml cefotaxime and the relevant plant resistance antibiotic (25 µg/ml hygromycin or 80 µg/ml kanamycin), which were incubated in *in vitro* growth conditions. The explants were transferred to fresh plates every 2 weeks until calli were formed, after which one callus per explant was removed and transferred to fresh plates every 2 weeks until reaching 1-2 cm in diameter. At this point, calli were transferred to MS2 medium (MS1 without MES or TDZ, with 3.5 g/L gelrite instead of agar) at pH 5.6 supplemented with 500 µg/ml cefotaxime and incubated in *in vitro* conditions until shoots were formed. Finally, shoot cuttings were transferred to *in vitro* growth medium and maintained in *in vitro* conditions.

### Selection of transformed lines

*ProMYB305a:GFP* lines were screened by confocal laser scanning microscopy on the adaxial epidermis of the terminal leaves. Lines in which GFP fluorescence was visible were selected for further imaging and ultimately, the line with the strongest GFP fluorescence was used for further experiments. CRISPR/Cas9-generated mutant lines of *MYB305a* (Potrx042905g12730) and *MYB305b* (Potrx058201g19640) in T89 were genotyped with Phire Plant Direct PCR Master Mix (Thermo Scientific) according to the manufacturer’s instructions, using the primers PttMYB305a_2sgR_gF/R and PttMYB305b_4sgR_gF/R (Table S1). The PCR product sizes suggested that the expected large-fragment deletions were generated for *MYB305a* but not for *MYB305b*. Products were extracted from the PCR gels using the QIAquick Gel Extraction Kit (Qiagen), cloned into chemically competent TOP10 *Escherichia coli* with the Zero Blunt TOPO PCR Cloning Kit (Invitrogen), and purified from *E. coli* transformants selected on 50 µg/ml kanamycin using the QIAprep Spin Miniprep Kit (Qiagen), according to the instructions of the kit manufacturers. The amplified plasmids were then sent to Eurofins Genomics for sequencing to determine actual DNA sequence alterations. CLC Main Workbench (Qiagen) software was used for sequence alignments. For further experiments, lines of *myb305a* were selected in which both alleles of *MYB305a* contained a large-fragment deletion at the expected site. Lines of *myb305ab* were selected in which both alleles of *MYB305a* and *MYB305b* contained either a large-fragment deletion, a large-fragment inversion, or indels that were expected to result in a frameshift mutation, at the expected sites. *Pro35S:MYB305a* lines were screened by RT-qPCR analysis of *MYB305a* expression levels in leaf samples from *in vitro* plants. Lines in which gene expression was enhanced compared to T89 were then grown in soil and analyzed again by RT-qPCR for *MYB305a* expression levels in terminal leaves. Those lines in which gene expression was enhanced compared to T89 (2 to 10,000-fold), without affecting plant height, were selected for further experiments.

### RT-qPCR

RNA was extracted using the RNeasy Plant Mini Kit (Qiagen) according to the manufacturer’s instructions. Leaf blade samples, excluding the petiole and mid vein, were ground in liquid nitrogen using a pestle and mortar and no more than 100 mg frozen ground tissue was used per sample for RNA extraction. The spin column membrane washing step was split into two washes of half the suggested buffer volume, with a 15-min treatment with 80 µl RQ1 DNase (Promega) in between the washes. RNA concentrations were measured with a Nano-Drop and cDNA was prepared from 1 µg RNA using the iScript cDNA Synthesis Kit (Bio-Rad) according to the manufacturer’s instructions. PCR was performed with cDNA using DreamTaq reagents (Thermo Scientific) and the primers PttMYB305a_RT-qPCR_F1/R1 for visualization of amplified products in an agarose gel. RT-qPCR was performed on a CFX96 Touch real-time PCR detection system (Bio-Rad) using Sso Advanced SYBR Green Supermix (Bio-Rad) and the primers PttMYB305a_RT-qPCR_F2/R2 for *MYB305a* (Potra2n8c18226) in European aspen, PttMYB305a_RT-qPCR_F1/R1 for *MYB305a* (Potrx042905g12730) in T89 and PtUBQ_qRT-PCR_F/R for the *UBIQUITIN* house-keeping gene (designed for Potra2n1c3634 in European aspen). See Table S1 for primer sequences. The 2^-ΔΔCt^ method^66^ was used to calculate relative gene expression.

### *k*-means clustering and variance partitioning

The SwAsp individuals were clustered based on all cell shape and size features with *k-*means cluster analysis using the kmeans function in R with nstart = 25. The data were first tested with the silhouette, gap statistic (clusGap) and elbow methods in the ‘cluster’ package in R^67^ to determine the optimal number of clusters. We used genotype median values of published foliar carbon isotope discrimination data^23^ as a proxy water use efficiency indicator for the SwAsp genotypes. The bioclimatic variables ‘mean annual precipitation’ and ‘precipitation of the driest month’ averaged for the years 1970-2000 were extracted from WorldClim^68^ at 2.5 min resolution. Cluster membership assigned to SwAsp genotypes from the *k-*means cluster analysis was used as a factor to partition the variance for carbon isotope discrimination, as well as the two environmental variables, the latitude and the longitude at the original sampling sites of the SwAsp genotypes with separate tests for each variable.

## Author contributions

S.R. designed the research project. S.L., S.M.D., K.M.R. and Z.R. performed the research and analyzed the data. All authors interpreted the data. S.R. and N.R.S. provided project supervision and contributed new tools. S.M.D. wrote the paper with input from all authors.

## Acknowledgements

We thank Skogforsk for hosting the SwAsp common garden experiment, Peter Grones, Aisha Beshirova, Filip Buchel and Anna Oravetzová for technical assistance and the Knut and Alice Wallenberg Foundation for essential financing of Umeå Plant Science Centre via Wallenberg Initiatives in Forest Research (WIFORCE). We are grateful to the UPSC Microscopy Facility for access to imaging equipment, the UPSC Poplar Transgenics Facility for *Populus* transformation and *in vitro* propagation and the UPSC Tree Phenotyping Platform for access to the automated conveyor network and controlled growth room and we thank the technical staff at these facilities and the “Green Team” at the UPSC Growth Facilities for their assistance. We acknowledge the Swedish Research Council (S.M.D., S.R., grant VR-2020-03420 to S.L.), the Kempe Foundations (grant JCK-1732 to S.L., grant JCK-1912 to Z.R.), the Knut and Alice Wallenberg Foundation (S.L.), VINNOVA (S.M.D., S.R.), the European Research Council (grant ERC-2024-SyG STARMORPH 101166880 to S.M.D.) and the Trees and Crops for the Future (TC4F) Strategic Research Area funding (K.M.R, N.R.S.) for funding this work.

## Supplementary Material

**Supplementary Fig. S1: Pavement cell features of representative Swedish aspen genotypes of more and less complex cell shape in the field.** A-D, circularity (A), branch count (B), lobe number (C) and area (D) of pavement cells in adaxial epidermis of pre-formed leaf in adult trees of representative SwAsp European aspen genotypes of more and less complex cell shape grown in the field. Each data point in the box plot represents the value of one cell; 70-160 cells per leaf, 2-3 leaves per tree and 1-5 trees per genotype were analyzed. Different letters indicate significantly different distributions at *P* < 0.05 (Wilcoxon rank sum test).

**Supplementary Fig. S2: Pavement cell images of representative Swedish aspen genotypes of more and less complex cell shape in the field.** A-H, representative confocal microscopy images of fluorescence in calcofluor-stained pavement cells in adaxial epidermis of pre-formed leaf in adult trees from the SwAsp collection of European aspen genotypes grown in the field. Maximum projections of z-stacks are shown. In the right panels, free-hand cell outline drawings are overlayed onto the images shown in the left panels. Those genotypes selected as representatives of more and less complex cell shape are shown in A-D (SwAsp numbers in magenta) and E-H (SwAsp numbers in cyan), respectively. Scale bars represent 50 μm.

**Supplementary Fig. S3: Identification of a SNP in *MYB305a* associated with pavement cell shape complexity in aspen.** A, quantile-quantile (QQ) plot of the *P*-values obtained for branch count in the GWAS. B-J, circularity (B), lobe number (C), solidity (D), lobeyness (E), non-lobe area (F), cell area (G), perimeter (H), length (I) and width (J) of pavement cells in the SwAsp genotypes analyzed, grouped according to the alleles of the candidate SNP in *MYB305a* present in the genotypes.

**Supplementary Fig. S4: Broad-sense heritability and genetic correlations of the pavement cell features in aspen.** A-B, broad-sense heritability (A) and heat map of genetic correlations (B) of the shape- and size-related pavement cell features analyzed in the SwAsp genotypes.

**Supplementary Fig. S5: Expression of *MYB305a* in Swedish aspen genotypes and the closest phylogenetic relatives of *MYB305a* in aspen and Arabidopsis.** A, genomic sequence of *MYB305a* in European aspen indicating the annealing sites of two primer pairs used to analyze gene expression, one of which spans the candidate T/C SNP (cyan arrows) and the other of which does not (magenta arrows). Introns are shown as black lines, UTRs as red lines and exons as cyan boxes (A). B-C, expression of *MYB305a* in the representative European aspen SwAsp genotypes for more and less complex pavement cell shape (indicated in magenta and cyan, respectively), as analyzed by visualization of cDNA PCR products amplified using the primer pair spanning the candidate SNP (B), and by RT-qPCR analysis using the primer pair not spanning the candidate SNP (C). Products for the house-keeping gene *UBIQUITIN* (*UBQ*) are also shown in B. Expression of *MYB305a* relative to that in SwAsp036 is shown in C; error bars represent standard error of the mean. D, phylogenetic relationships of the closest homologs of *MYB305a* in European aspen and Arabidopsis, made using the pre-generated phylogenetic trees constructed as part of a large *Picea abies* (Norway spruce) and *Pinus sylvestris* (Scots pine) genome project^69^.

**Supplementary Fig. S6: Pavement cell shape complexity varies among genotypes of juvenile aspen in the greenhouse.** A-H, representative light microscopy images of pavement cells in adaxial epidermis of terminal leaf number 17 in juvenile trees from the SwAsp collection of European aspen genotypes grown in the greenhouse. In the right panels, free-hand cell outline drawings of cells in the left panel images are shown. Those genotypes selected as representatives of more and less complex cell shape are shown in A-D (SwAsp numbers in magenta) and E-H (SwAsp numbers in cyan), respectively. Scale bars represent 25 μm.

**Supplementary Fig. S7: Pavement cell shape complexity in terminal leaves of juvenile hybrid aspen clone T89 in the greenhouse.** A-D, circularity (A), branch count (B), lobe number (C) and area (D) of pavement cells in adaxial epidermis of terminal leaf numbers 7-19 in hybrid aspen clone T89 grown in the greenhouse. Leaf number was counted from the top down, with number 1 being the youngest leaf. Each data point represents the mean value of 89-605 cells in total, measured across 1 leaf per leaf number per tree and 3 trees. Error bars represent standard error of the mean.

**Supplementary Fig. S8: Selection of genetically transformed *ProMYB305a:GFP*, *MYB305a* CRISPR/Cas9 deletion and *Pro35S:MYB305a* over-expressor lines of hybrid aspen (T89).** A, representative confocal microscopy images of GFP fluorescence in pavement cells in adaxial epidermis of terminal leaf number 12 in 3 lines of *ProMYB305a:GFP* grown in the greenhouse. Due to weak fluorescence, image acquisition settings were varied among the lines. Scale bars represent 25 μm. B, guide RNA (gRNA) target sites (magenta arrowheads) in the genomic and coding sequences of *MYB305a* (Potrx042905g12730) in T89. Introns are shown as black lines and exons as cyan boxes. C, genomic sequences of *MYB305a* in T89 and 6 *MYB305a* CRISPR/Cas9 large-fragment deletion lines (*myb305a*) at the sites of expected deletion, showing gRNA sequences (in magenta), PAM sequences (in cyan), sites of expected DNA break (magenta arrowheads) and actual deletions obtained (red dashes represent missing nucleotides). For simplicity, the genomic sequence between the gRNAs is not shown (represented as dotted lines). Two sequences are shown for those lines for which different deletions were obtained in each allele. The obtained deletion sizes are shown to the right. D, expression of *MYB305a* relative to that in T89 in terminal leaf number 17 in 5 *MYB305a* over-expressor (*Pro35S:MYB305a*, in magenta) lines grown in the greenhouse. Error bars represent standard error of the mean. Different letters indicate significantly different means at *P* < 0.05 (Tukey’s HSD test).

**Supplementary Fig. S9: Shoot height in *MYB305a* CRISPR/Cas9 deletion and *Pro35S:MYB305a* over-expressor lines of hybrid aspen (T89).** A-D, shoot height in *myb305a* mutant lines (A-B) and *Pro35S:MYB305a* over-expressor lines (C-D) compared to T89 after 8 weeks of growth in soil. Images of representative plants are shown; side shoots were removed 3 and 6 weeks after potting in soil; scale bars represent 30 cm. Error bars represent standard error of the mean. Different letters indicate significantly different means at *P* < 0.05 (Tukey’s HSD test; NS, not significant).

**Supplementary Fig. S10: Line selection and shoot height in *MYB305a MYB305b* CRISPR/Cas9 deletion lines of hybrid aspen (T89).** A, genomic sequences of *MYB305a* in T89 and 11 *MYB305a MYB305b* CRISPR/Cas9 mutant (*myb305ab*) lines at the sites of expected deletion, showing guide RNA (gRNA) sequences (in magenta), PAM sequences (in cyan), sites of expected DNA break (magenta arrowheads) and actual deletions (red dashes represent missing nucleotides) or insertions (in red) obtained. An inversion was discovered in two of the lines (black boxes). For simplicity, the genomic sequence between the gRNAs is not shown (represented as dotted lines). Two sequences are shown for those lines for which different deletions were obtained in each allele. The obtained deletion/insertion sizes are shown to the right. B, gRNA target sites (magenta arrowheads) in the genomic and coding sequences of *MYB305b* (Potrx058201g19640) in T89. Introns are shown as black lines and exons as cyan boxes. C, genomic sequences of *MYB305b* in T89 and the same 11 *myb305ab* lines for which *MYB305a* is shown in (A). D-E, shoot height in *myb305ab* mutant lines compared to T89 after 8 weeks of growth in soil. Side shoots were removed 3 and 6 weeks after potting in soil; scale bars represent 30 cm. Error bars represent standard error of the mean. NS, not significantly different means at *P* < 0.05 (Tukey’s HSD test).

**Supplementary Fig. S11: Pavement cell shape features in *MYB305a MYB305b* CRISPR/Cas9 deletion lines.** A-D, circularity (A), branch count (B), lobe number (C) and area (D) of pavement cells in adaxial epidermis of terminal leaf number 17 in double *MYB305a MYB305b* CRISPR mutant (*myb305ab*, in cyan) lines of hybrid aspen (T89) grown in the greenhouse. Each data point in the box plot represents one of 118-145 cells in total measured across 1 leaf per tree and 3 trees per genotype. Different letters indicate significantly different distributions or means at *P* < 0.05 (Wilcoxon rank sum test for A-C; Tukey’s HSD test for D).

**Supplementary Fig. S12: Effects of drought stress on shoot height and pavement cell shape features in hybrid aspen (T89) leaves of different developmental stages.** A-B, shoot height in *ProMYB305a:GFP* line 4 in control and drought conditions at time of leaf sampling for pavement cell imaging (after 7 weeks of growth in soil). Images of representative plants are shown; side shoots were removed 3 and 6 weeks after potting in soil; scale bar represents 20 cm. Error bars represent standard error of the mean. Asterisks indicate significantly different means between the control and drought treatment according to the Student’s t-test (\*\*\**P* < 0.001). C-F, circularity (C), branch count (D), lobe number (E) and area (F) of pavement cells in adaxial epidermis of terminal leaf numbers 9-19 in *ProMYB305a:GFP* line 4 of hybrid aspen (T89) grown in control or drought conditions in the greenhouse (see Fig. 6A-F for representative images). Each data point in the box plot represents one of 156-485 cells in total measured across 1 leaf per developmental stage per tree and 3 trees per treatment. Asterisks indicate significantly different distributions between the control and drought treatment according to the Wilcoxon rank sum test (NS, not significant; \**P* < 0.05; \*\**P* < 0.01; \*\*\**P* < 0.001).

**Supplementary Fig. S13: Effects of drought stress on shoot height in *MYB305a* CRISPR/Cas9 deletion and *Pro35S:MYB305a* over-expressor lines of hybrid aspen (T89).** A-B, shoot height in T89, *myb305a* mutant lines and *Pro35S:MYB305a* over-expressor lines in control and drought conditions at time of leaf sampling for pavement cell imaging (after 7 weeks of growth in soil). Images of representative plants are shown; side shoots were removed 3 and 6 weeks after potting in soil; scale bars represent 30 cm. Error bars represent standard error of the mean. Asterisks indicate significantly different means between the control and drought treatment according to the Student’s t-test (\**P* < 0.05; \*\**P* < 0.01; \*\*\**P* < 0.001).

**Supplementary Fig. S14: *k*-means clustering assigns the Swedish aspen genotypes into more and less complex pavement cell shape clusters.** A, *k*-means cluster analysis of the SwAsp genotypes based on all cell shape and size features. The representative genotypes for more and less complex pavement cell shape are indicated in magenta and cyan, respectively. B, branch of pavement cells in the SwAsp genotypes, as shown in Fig. 1D, but color-coded according to the cluster to which that genotype was assigned by *k*-means cluster analysis, with cyan used for cluster 1 and magenta for cluster 2.

**Supplementary Fig. S15: The cell feature clusters associate significantly with original sampling site environments.** A-E, cluster assignment of the SwAsp genotypes from the *k-*means cluster analysis was used as a factor to partition the variance for foliar carbon isotope discrimination (delta^13^C) (A), precipitation in the driest month of the year averaged over 30 years (B), annual precipitation averaged over 30 years (C), latitude (D) and longitude (E) at the original sampling sites. The *P*-values indicated were calculated using the Wilcoxon rank sum test.

**Supplementary Table S1: Primers used in this study.** Names and sequences of all primers used in this study.

**Supplementary Data S1: Pavement cell shape- and size-related features of Swedish aspen genotypes in the field.** Means and standard deviations were calculated from measurements of 70-160 cells per leaf, 2-3 leaves per tree and 1-5 trees per genotype.

**Supplementary Data S2: The top-ranked 1000 genes for each cell feature based on significance of the association of its SNPs with that feature.** The candidate T-to-C SNP in *MYB305a* (chr8_11748682_C_T) is indicated in red.

