## Supplementary Material for "A drought stress-induced MYB transcription factor regulates pavement cell shape in leaves of European aspen (*Populus tremula*)"

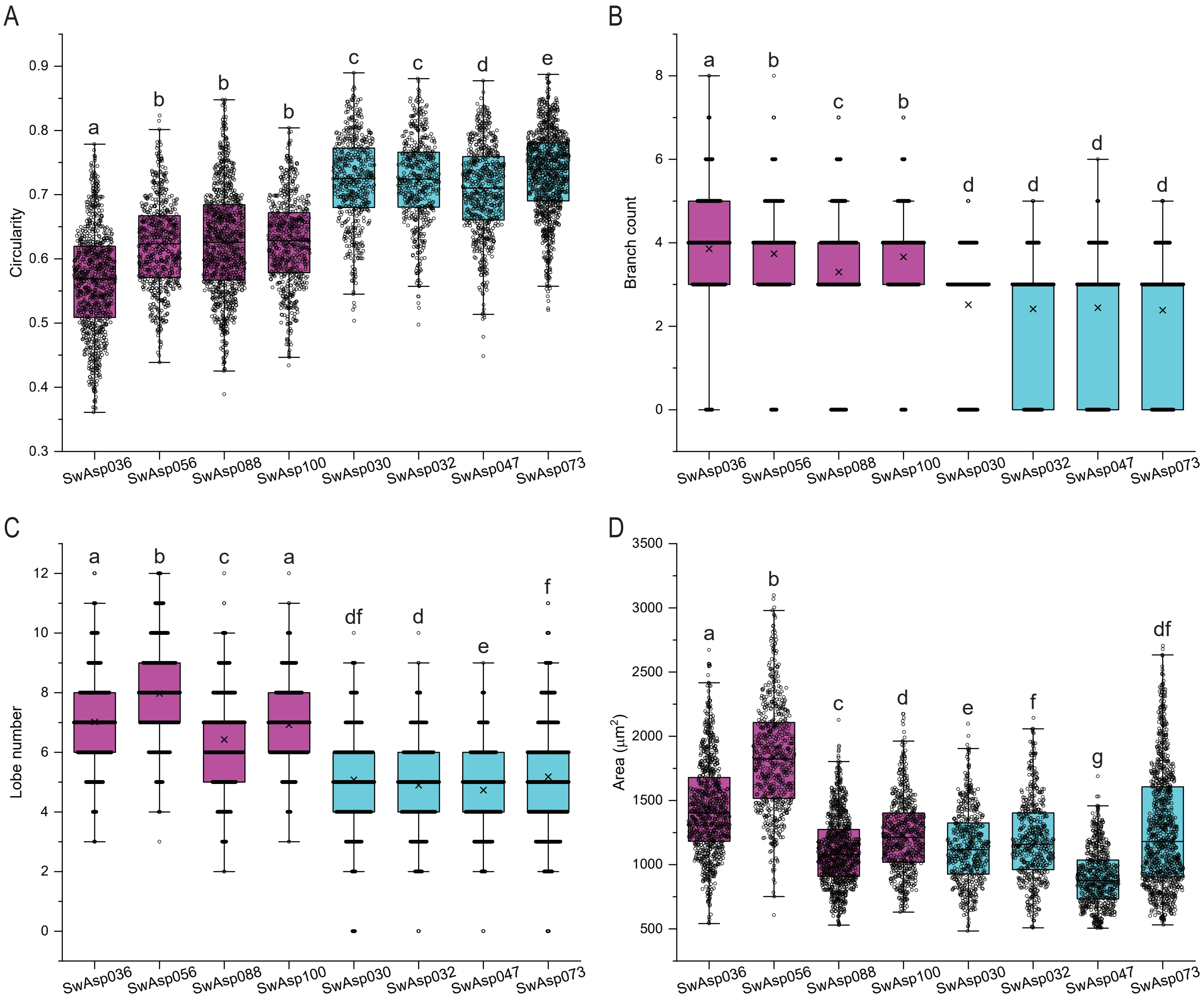


**Supplementary Fig. S1: Pavement cell features of representative Swedish aspen genotypes of more and less complex cell shape in the field.** A-D, circularity (A), branch count (B), lobe number (C) and area (D) of pavement cells in adaxial epidermis of pre-formed leaf in adult trees of representative SwAsp European aspen genotypes of more and less complex cell shape grown in the field. Each data point in the box plot represents the value of one cell; 70-160 cells per leaf, 2-3 leaves per tree and 1-5 trees per genotype were analyzed. Different letters indicate significantly different distributions at $P < 0.05$ (Wilcoxon rank sum test).

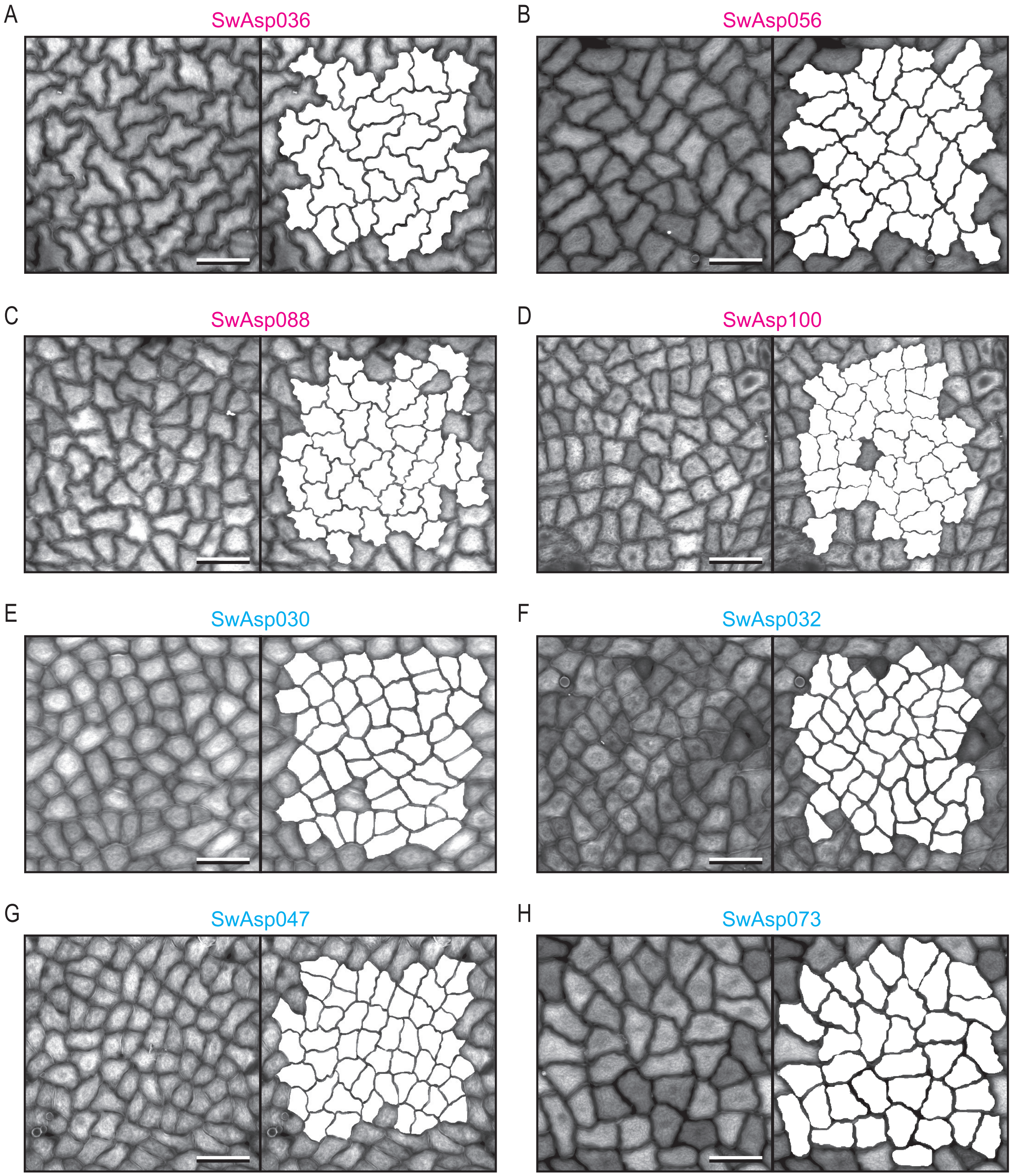


**Supplementary Fig. S2: Pavement cell images of representative Swedish aspen genotypes of more and less complex cell shape in the field.** A-H, representative confocal microscopy images of fluorescence in calcofluor-stained pavement cells in adaxial epidermis of pre-formed leaf in adult trees from the SwAsp collection of European aspen genotypes grown in the field. Maximum projections of z-stacks are shown. In the right panels, free-hand cell outline drawings are overlayed onto the images shown in the left panels. Those genotypes selected as representatives of more and less complex cell shape are shown in A-D (SwAsp numbers in magenta) and E-H (SwAsp numbers in cyan), respectively. Scale bars represent 50 µm.

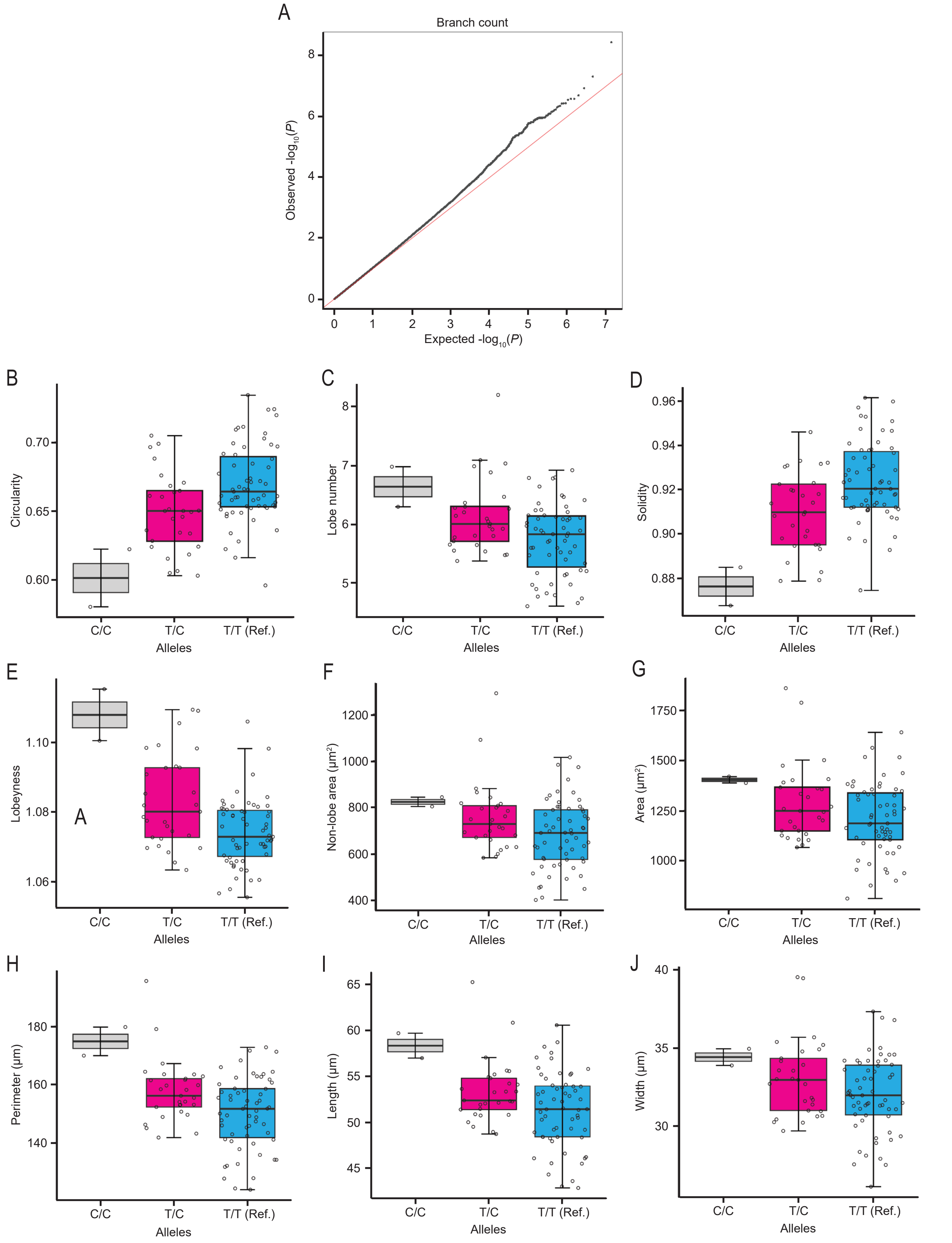


**Supplementary Fig. S3: Identification of a SNP in *MYB305a* associated with pavement cell shape complexity in aspen.** A, quantile-quantile (QQ) plot of the *P*-values obtained for branch count in the GWAS. B-J, circularity (B), lobe number (C), solidity (D), lobeyness (E), non-lobe area (F), cell area (G), perimeter (H), length (I) and width (J) of pavement cells in the SwAsp genotypes analyzed, grouped according to the alleles of the candidate SNP in *MYB305a* present in the genotypes.

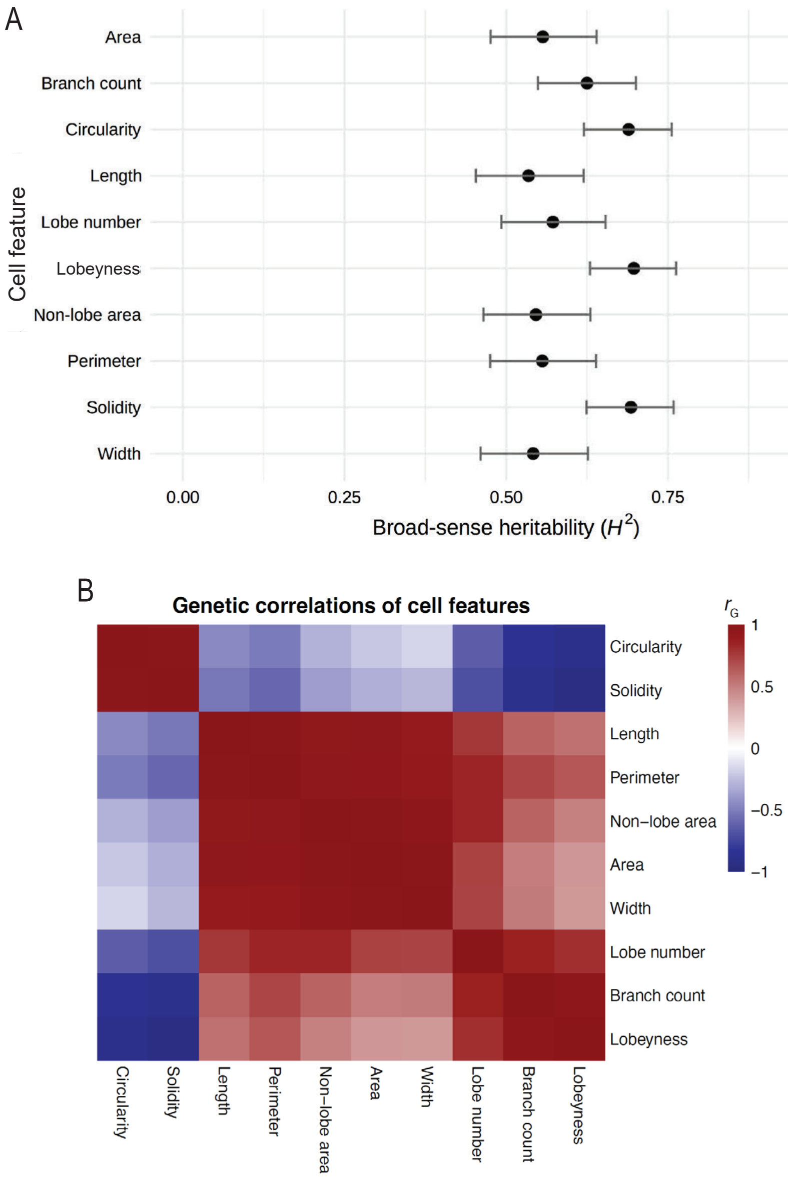


**Supplementary Fig. S4: Broad-sense heritability and genetic correlations of the pavement cell features in aspen.** A-B, broad-sense heritability (A) and heat map of genetic correlations (B) of the shape- and size-related pavement cell features analyzed in the SwAsp genotypes.

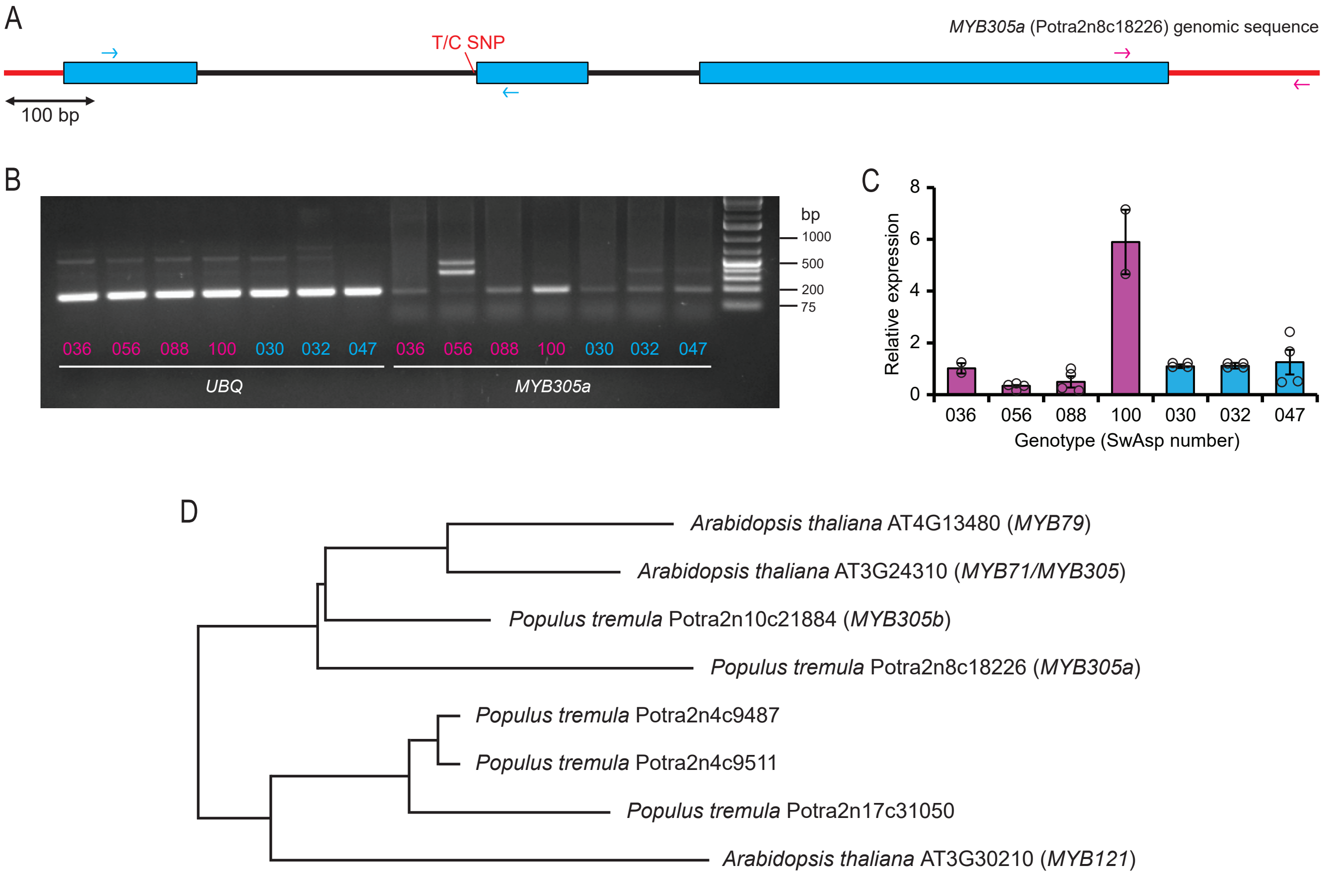


**Supplementary Fig. S5: Expression of *MYB305a* in Swedish aspen genotypes and the closest phylogenetic relatives of *MYB305a* in aspen and Arabidopsis.** A, genomic sequence of *MYB305a* in European aspen, indicating the annealing sites of two primer pairs used to analyze gene expression, one of which spans the candidate T/C SNP (cyan arrows) and the other of which does not (magenta arrows). Introns are shown as black lines, UTRs as red lines and exons as cyan boxes (A). B-C, expression of *MYB305a* in the representative European aspen SwAsp genotypes for more and less complex pavement cell shape (indicated in magenta and cyan, respectively), as analyzed by visualization of cDNA PCR products amplified using the primer pair spanning the candidate SNP (B), and by RT-qPCR analysis using the primer pair not spanning the candidate SNP (C). Products for the house-keeping gene *UBIQUITIN* (*UBQ*) are also shown in B. Expression of *MYB305a* relative to that in SwAsp036 is shown in C; error bars represent standard error of the mean. D, phylogenetic relationships of the closest homologs of *MYB305a* in European aspen and Arabidopsis, made using the pre-generated phylogenetic trees constructed as part of a large *Picea abies* (Norway spruce) and *Pinus sylvestris* (Scots pine) genome project[69].

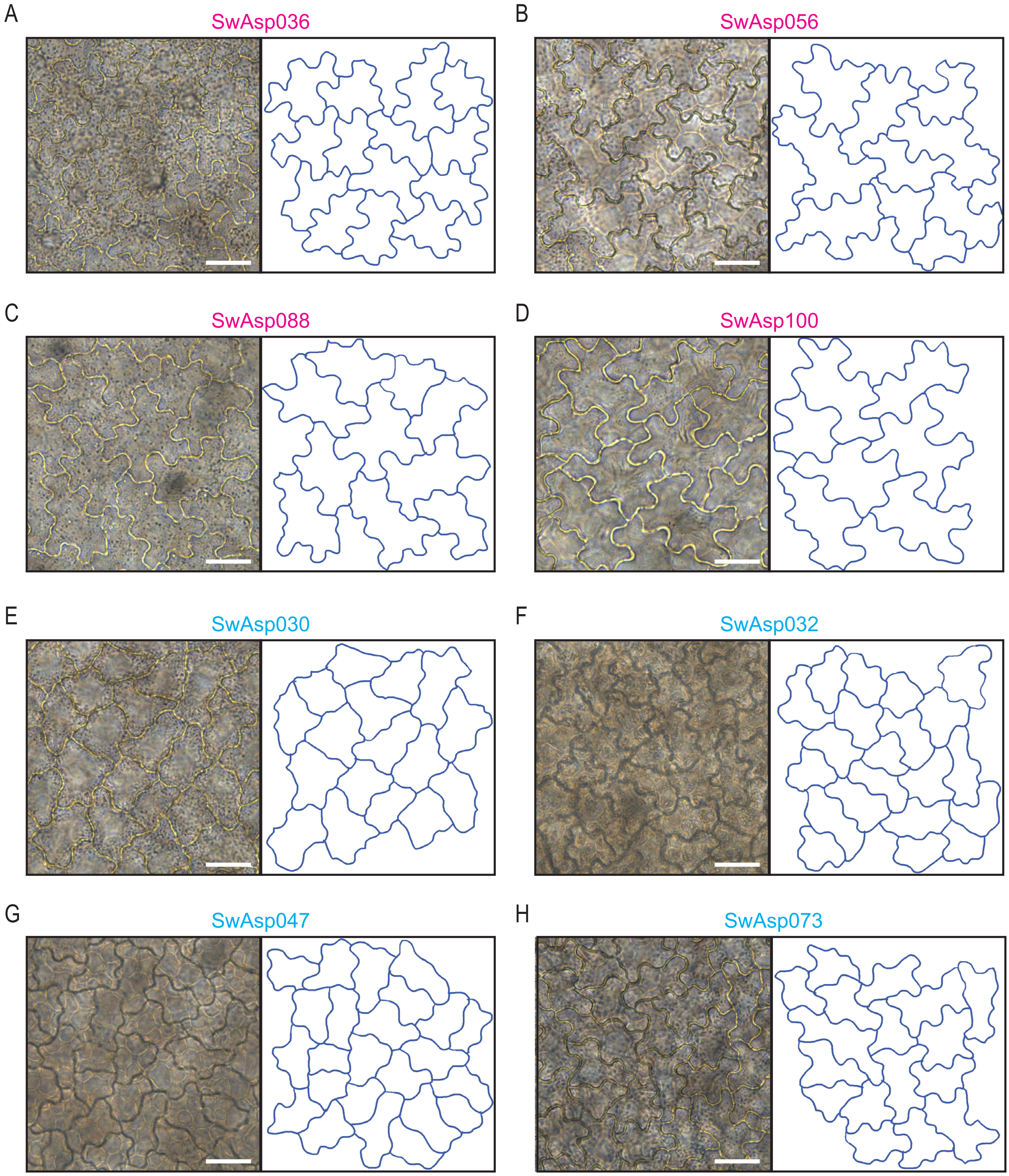


**Supplementary Fig. S6: Pavement cell shape complexity varies among genotypes of juvenile aspen in the greenhouse.** A-H, representative light microscopy images of pavement cells in adaxial epidermis of terminal leaf number 17 in juvenile trees from the SwAsp collection of European aspen genotypes grown in the greenhouse. In the right panels, free-hand cell outline drawings of cells in the left panel images are shown. Those genotypes selected as representatives of more and less complex cell shape are shown in A-D (SwAsp numbers in magenta) and E-H (SwAsp numbers in cyan), respectively. Scale bars represent 25 µm.

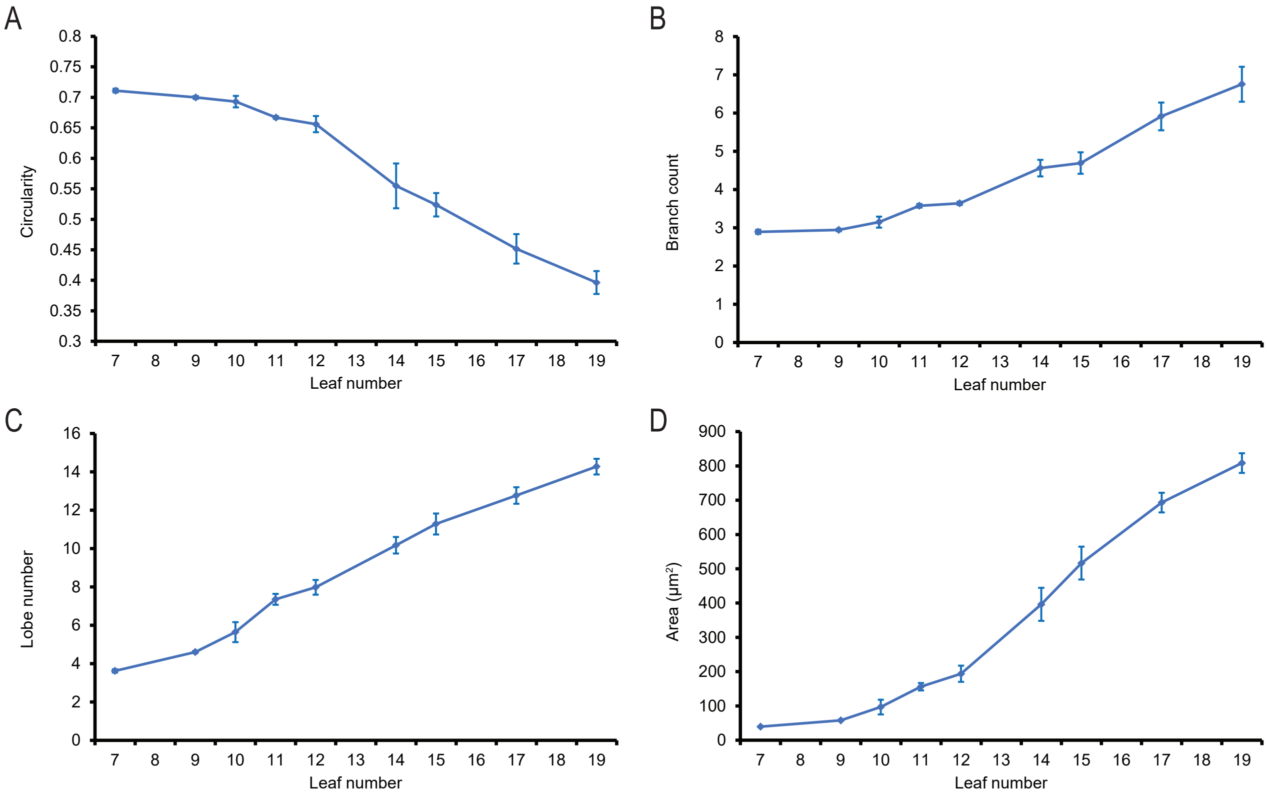


**Supplementary Fig. S7: Pavement cell shape complexity in terminal leaves of juvenile hybrid aspen clone T89 in the greenhouse.** A-D, circularity (A), branch count (B), lobe number (C) and area (D) of pavement cells in adaxial epidermis of terminal leaf numbers 7-19 in hybrid aspen clone T89 grown in the greenhouse. Leaf number counted from the top down, with number 1 being the youngest leaf. Each data point represents the mean value of 89--605 cells in total, measured across 1 leaf per leaf number per tree and 3 trees. Error bars represent standard error of the mean.

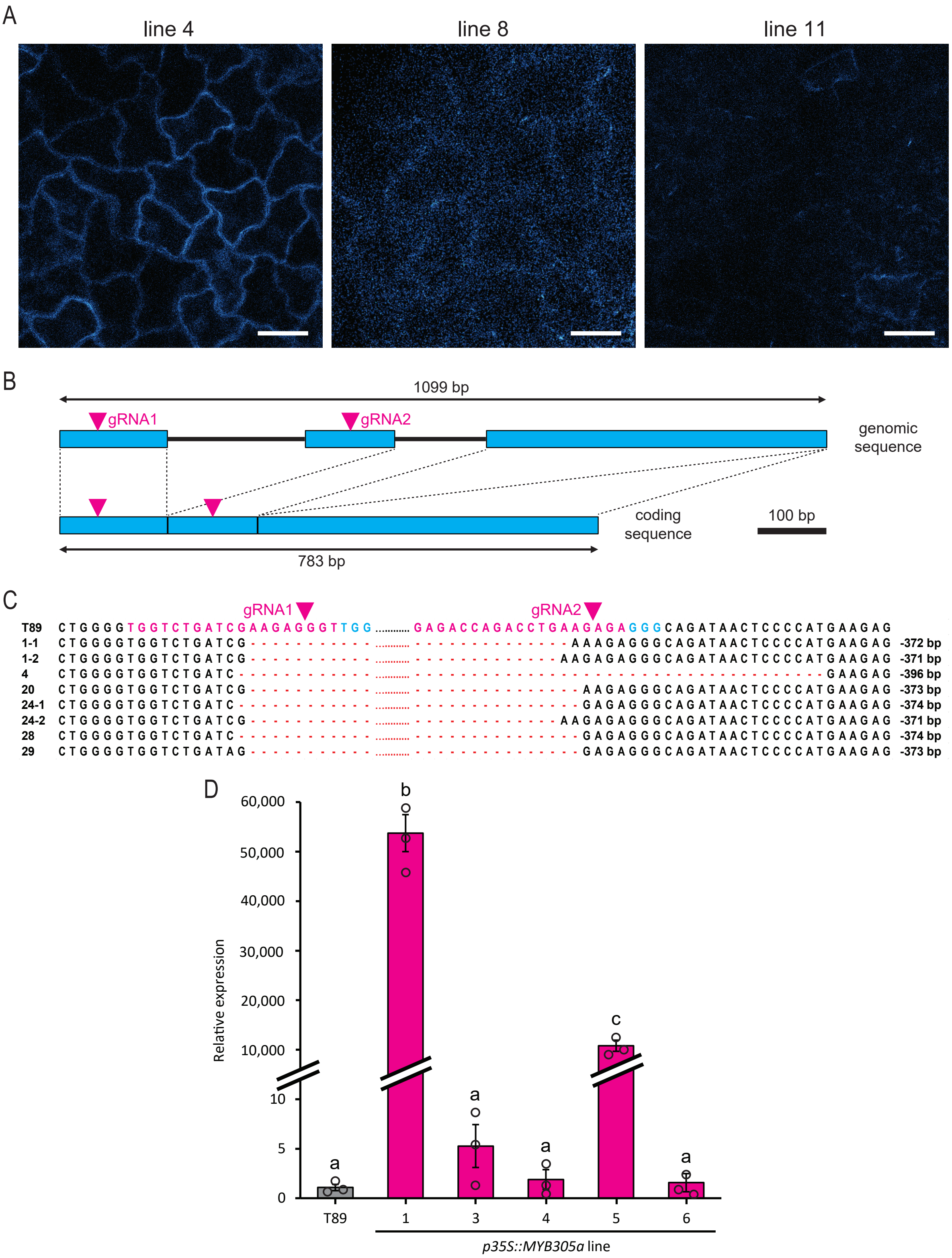


**Supplementary Fig. S8: Selection of genetically transformed *ProMYB305a:GFP*, *MYB305a* CRISPR/Cas9 deletion and *Pro35S:MYB305a* over-expressor lines of hybrid aspen (T89).** A, representative confocal microscopy images of GFP fluorescence in pavement cells in adaxial epidermis of terminal leaf number 12 in 3 lines of *ProMYB305a:GFP* grown in the greenhouse. Due to weak fluorescence, image acquisition settings were varied among the lines. Scale bars represent 25 µm. B, guide RNA (gRNA) target sites (magenta arrowheads) in the genomic and coding sequences of *MYB305a* (Potrx042905g12730) in T89. Introns are shown as black lines and exons as cyan boxes. C, genomic sequences of *MYB305a* in T89 and 6 *MYB305a* CRISPR/Cas9 large-fragment deletion lines (*myb305a*) at the sites of expected deletion, showing gRNA sequences (in magenta), PAM sequences (in cyan), sites of expected DNA break (magenta arrowheads) and actual deletions obtained (red dashes represent missing nucleotides). For simplicity, the genomic sequence between the gRNAs is not shown (represented as dotted lines). Two sequences are shown for those lines for which different deletions were obtained in each allele. The obtained deletion sizes are shown to the right. D, expression of *MYB305a* relative to that in T89 in terminal leaf number 17 in 5 *MYB305a* over-expressor (*Pro35S:MYB305a*, in magenta) lines grown in the greenhouse. Error bars represent standard error of the mean. Different letters indicate significantly different means at $P < 0.05$ (Tukey's HSD test).

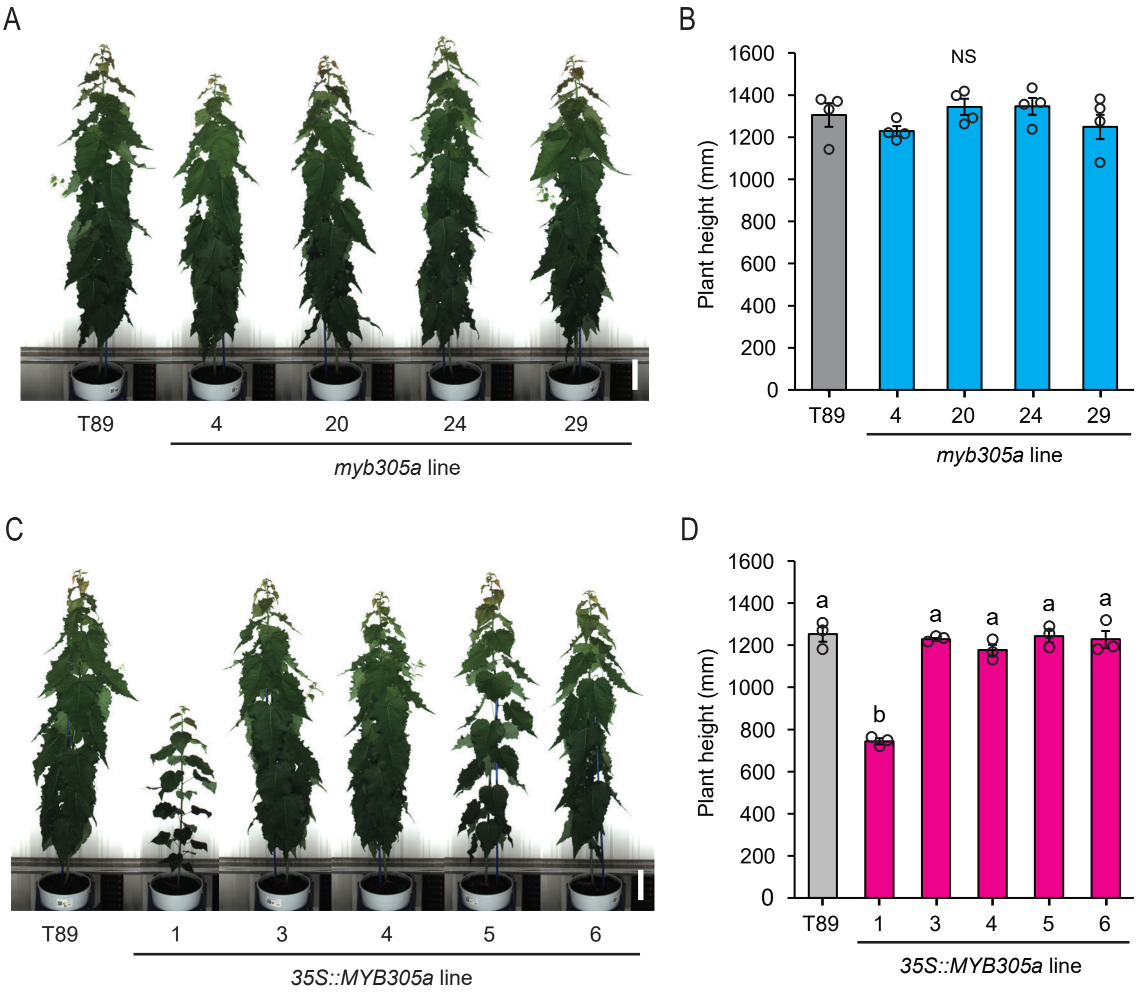


**Supplementary Fig. S9: Shoot height in *MYB305a* CRISPR/Cas9 deletion and *Pro35S:MYB305a* over-expressor lines of hybrid aspen (T89).** A-D, shoot height in *myb305a* mutant lines (A-B) and *Pro35S:MYB305a* over-expressor lines (C-D) compared to T89 after 8 weeks of growth in soil. Images of representative plants are shown; side shoots were removed 3 and 6 weeks after potting in soil; scale bars represent 30 cm. Error bars represent standard error of the mean. Different letters indicate significantly different means at $P < 0.05$ (Tukey's HSD test; NS, not significant).

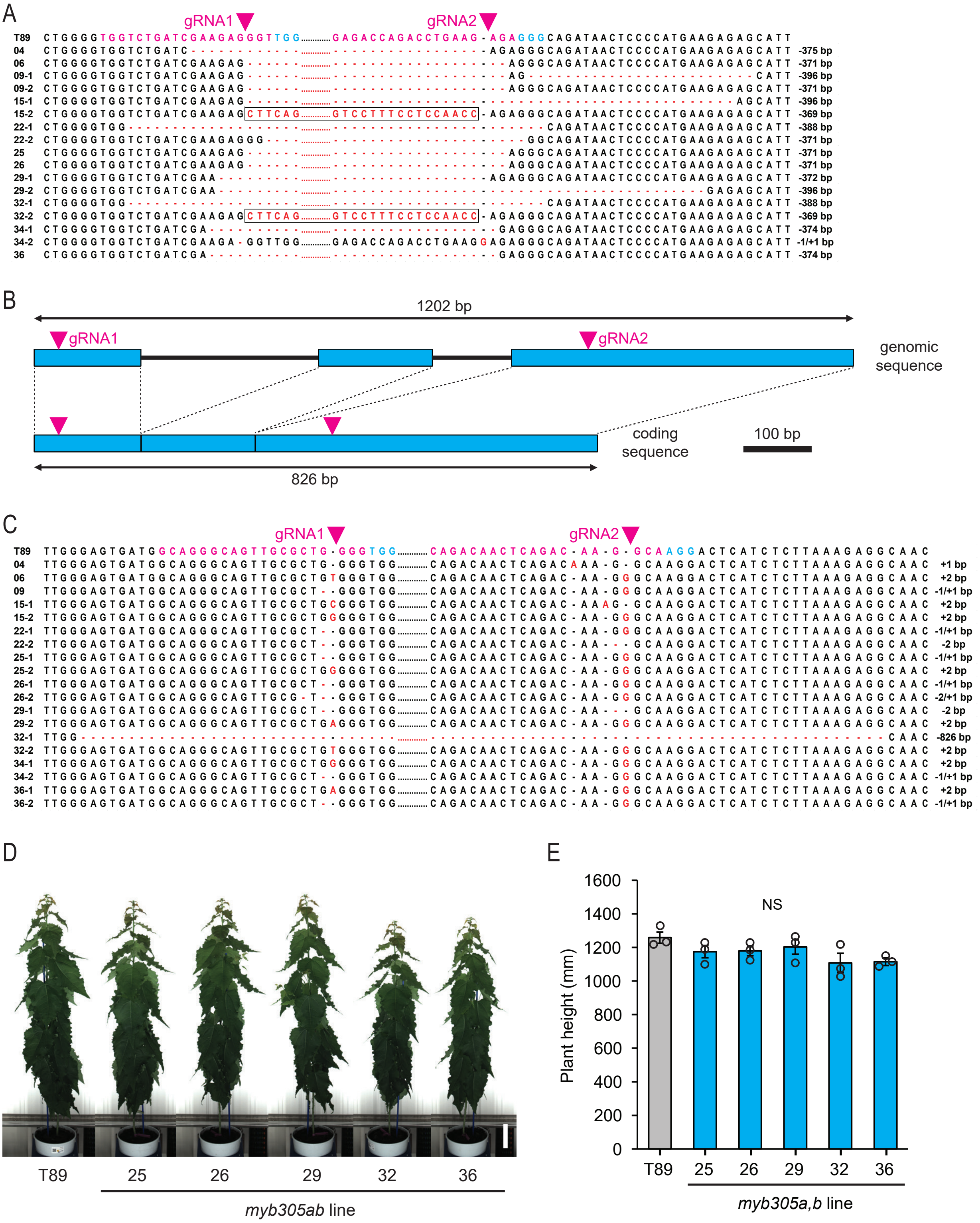


**Supplementary Fig. S10: Line selection and shoot height in *MYB305a MYB305b* CRISPR/Cas9 deletion lines of hybrid aspen (T89).** A, genomic sequences of *MYB305a* in T89 and 11 *MYB305a MYB305b* CRISPR/Cas9 mutant (*myb305ab*) lines at the sites of expected deletion, showing guide RNA (gRNA) sequences (in magenta), PAM sequences (in cyan), sites of expected DNA break (magenta arrowheads) and actual deletions (red dashes represent missing nucleotides) or insertions (in red) obtained. An inversion was discovered in two of the lines (black boxes). For simplicity, the genomic sequence between the gRNAs is not shown (represented as dotted lines). Two sequences are shown for those lines for which different deletions were obtained in each allele. The obtained deletion/insertion sizes are shown to the right. B, gRNA target sites (magenta arrowheads) in the genomic and coding sequences of *MYB305b* (Potrx058201g19640) in T89. Introns are shown as black lines and exons as cyan boxes. C, genomic sequences of *MYB305b* in T89 and the same 11 *myb305ab* lines for which *MYB305a* is shown in (A). D-E, shoot height in *myb305ab* mutant lines compared to T89 after 8 weeks of growth in soil. Side shoots were removed 3 and 6 weeks after potting in soil; scale bars represent 30 cm. Error bars represent standard error of the mean. NS, not significantly different means at $P < 0.05$ (Tukey's HSD test).

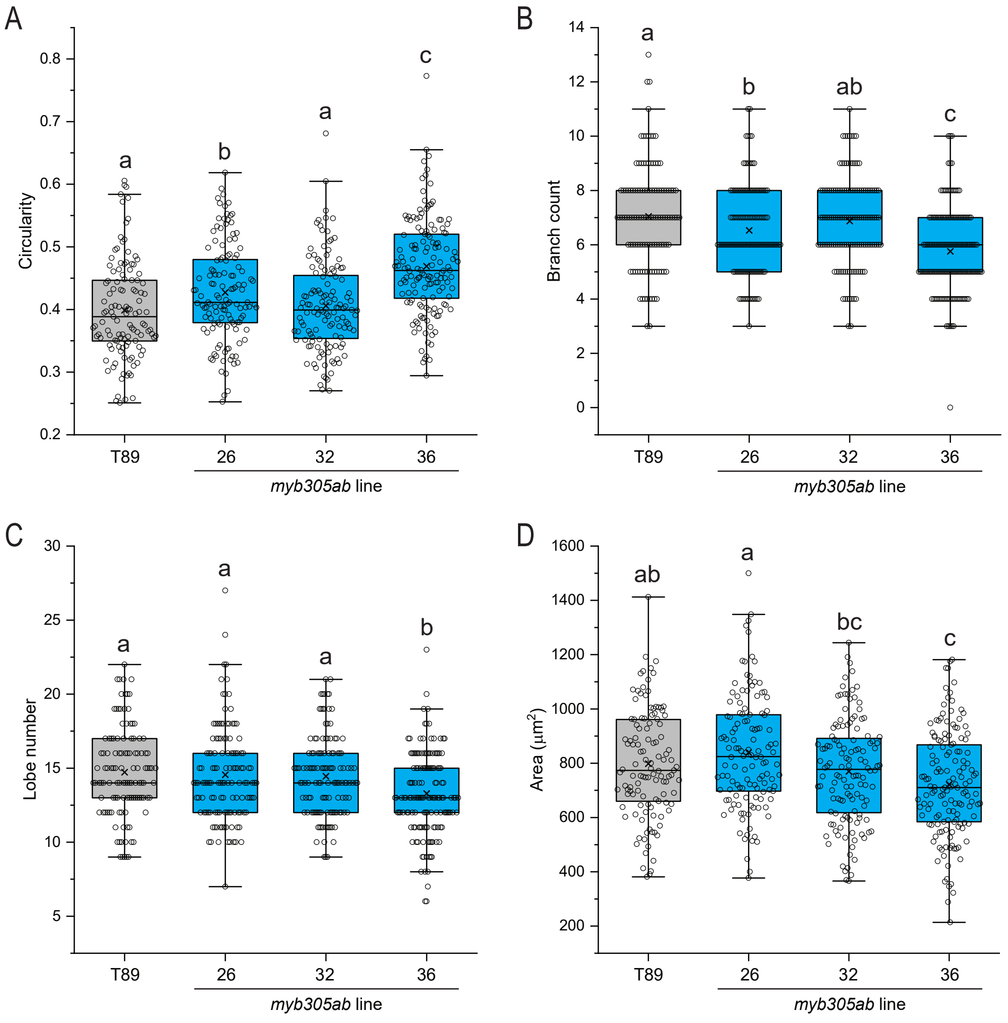


**Supplementary Fig. S11: Pavement cell shape features in *MYB305a MYB305b* CRISPR/Cas9 deletion lines.** A-D, circularity (A), branch count (B), lobe number (C) and area (D) of pavement cells in adaxial epidermis of terminal leaf number 17 in double *MYB305a MYB305b* CRISPR mutant (*myb305ab*, in cyan) lines of hybrid aspen (T89) grown in the greenhouse. Each data point in the box plot represents one of 118-145 cells in total measured across 1 leaf per tree and 3 trees per genotype. Different letters indicate significantly different distributions or means at $P < 0.05$ (Wilcoxon rank sum test for A-C; Tukey's HSD test for D).

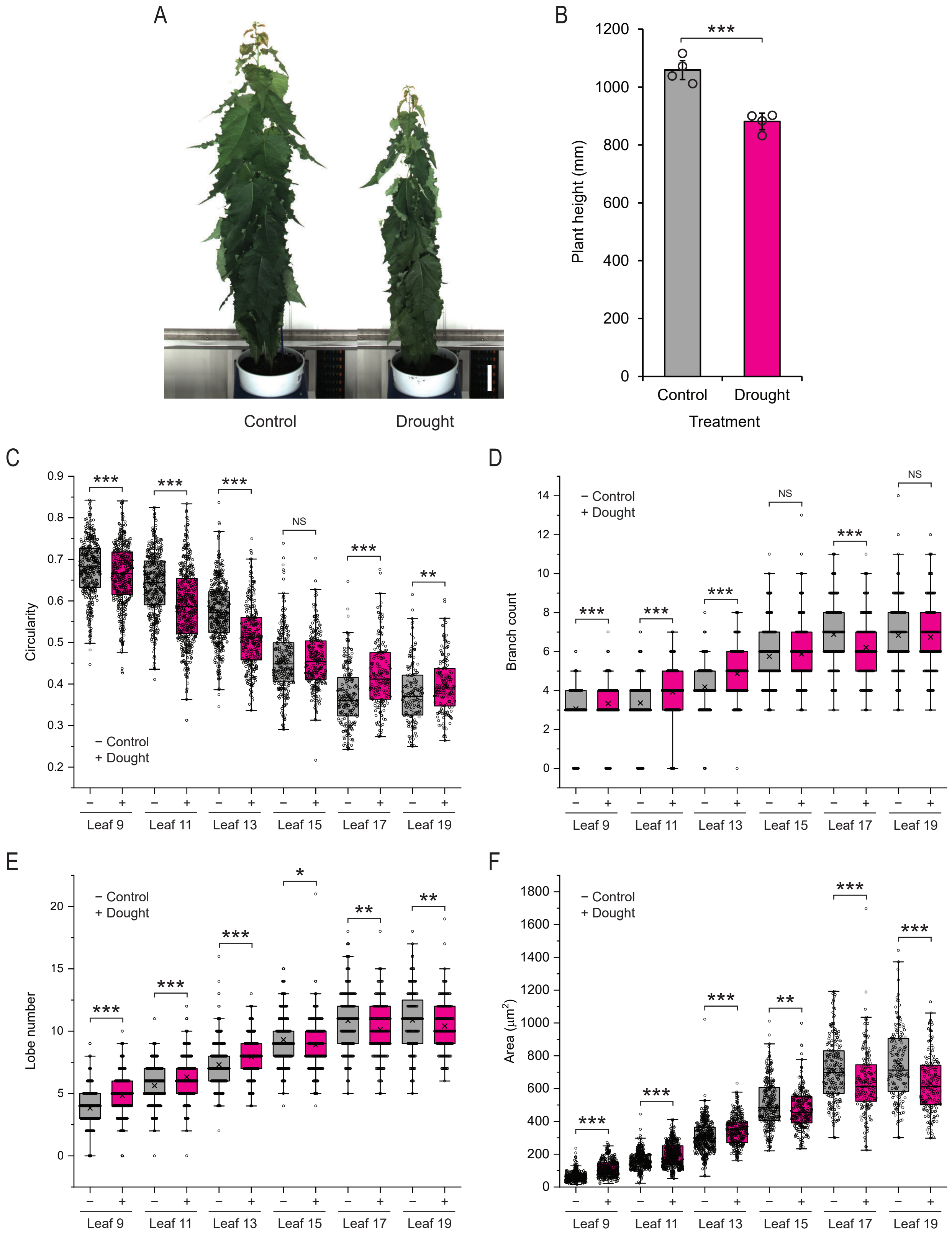


**Supplementary Fig. S12: Effects of drought stress on shoot height and pavement cell shape features in hybrid aspen (T89) leaves of different developmental stages.** A-B, shoot height in *ProMYB305a:GFP* line 4 in control and drought conditions at time of leaf sampling for pavement cell imaging (after 7 weeks of growth in soil). Images of representative plants are shown; side shoots were removed 3 and 6 weeks after potting in soil; scale bar represents 20 cm. Error bars represent standard error of the mean. Asterisks indicate significantly different means between the control and drought treatment according to the Student's t-test (****P* < 0.001). C-F, circularity (C), branch count (D), lobe number (E) and area (F) of pavement cells in adaxial epidermis of terminal leaf numbers 9-19 in *ProMYB305a:GFP* line 4 of hybrid aspen (T89) grown in control or drought conditions in the greenhouse (see Fig. 6A-F for representative images). Each data point in the box plot represents one of 156-485 cells in total measured across 1 leaf per developmental stage per tree and 3 trees per treatment. Asterisks indicate significantly different distributions between the control and drought treatment according to the Wilcoxon rank sum test (NS, not significant; **P* < 0.05; ***P* < 0.01; ****P* < 0.001).

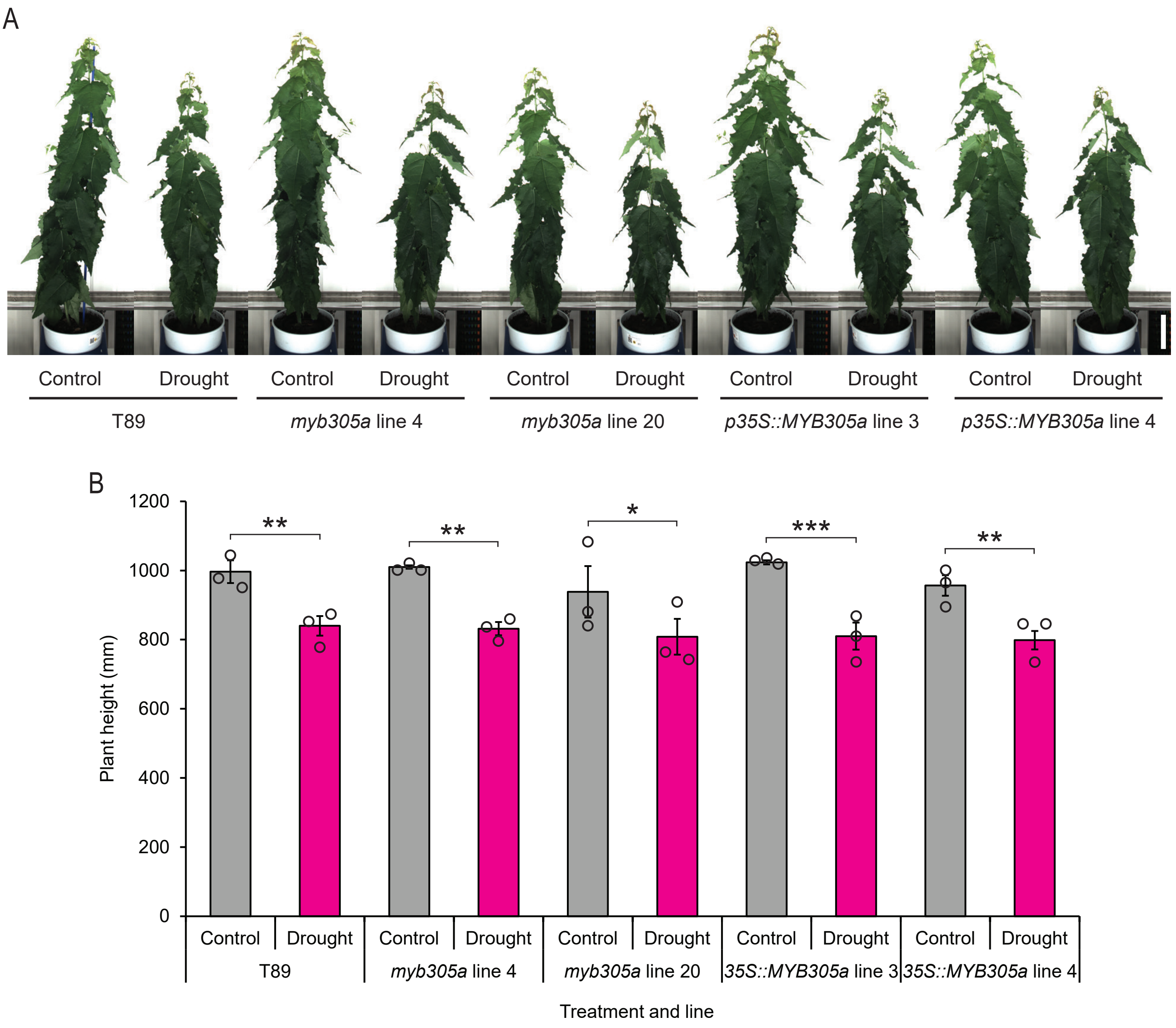


**Supplementary Fig. S13: Effects of drought stress on shoot height in *MYB305a* CRISPR/Cas9 deletion and *Pro35S:MYB305a* over-expressor lines of hybrid aspen (T89).** A-B, shoot height in T89, *myb305a* mutant lines and *Pro35S:MYB305a* over-expressor lines in control and drought conditions at time of leaf sampling for pavement cell imaging (after 7 weeks of growth in soil). Images of representative plants are shown; side shoots were removed 3 and 6 weeks after potting in soil; scale bars represent 30 cm. Error bars represent standard error of the mean. Asterisks indicate significantly different means between the control and drought treatment according to the Student's t-test (*$P$ < 0.05; **$P$ < 0.01; ***$P$ < 0.001).

A

| SwAsp no. | Cluster | SwAsp no. | Cluster | SwAsp no. | Cluster | SwAsp no. | Cluster |
|---|---|---|---|---|---|---|---|
| 001 | 2 | 034 | 2 | 061 | 2 | 088 | 2 |
| 002 | 1 | 035 | 2 | 062 | 1 | 089 | 2 |
| 003 | 2 | 036 | 2 | 063 | 1 | 090 | 2 |
| 004 | 1 | 037 | 1 | 064 | 2 | 091 | 2 |
| 005 | 1 | 038 | 2 | 065 | 1 | 092 | 1 |
| 006 | 2 | 039 | 1 | 066 | 2 | 093 | 1 |
| 007 | 2 | 040 | 2 | 067 | 1 | 094 | 2 |
| 009 | 1 | 041 | 1 | 068 | 1 | 095 | 2 |
| 011 | 1 | 042 | 2 | 069 | 1 | 096 | 2 |
| 012 | 2 | 043 | 1 | 070 | 1 | 097 | 2 |
| 013 | 2 | 044 | 2 | 071 | 1 | 098 | 1 |
| 014 | 1 | 045 | 2 | 072 | 1 | 099 | 2 |
| 018 | 1 | 046 | 2 | 073 | 1 | 100 | 2 |
| 019 | 2 | 047 | 1 | 074 | 1 | 103 | 2 |
| 020 | 1 | 048 | 2 | 075 | 2 | 104 | 2 |
| 022 | 2 | 049 | 1 | 076 | 2 | 105 | 1 |
| 023 | 1 | 050 | 1 | 077 | 2 | 106 | 2 |
| 024 | 1 | 051 | 2 | 078 | 2 | 108 | 2 |
| 025 | 1 | 052 | 2 | 079 | 1 | 110 | 2 |
| 026 | 2 | 053 | 2 | 080 | 2 | 111 | 2 |
| 028 | 1 | 055 | 1 | 081 | 1 | 112 | 2 |
| 029 | 1 | 056 | 2 | 082 | 1 | 113 | 2 |
| 030 | 1 | 057 | 2 | 084 | 2 | 114 | 2 |
| 031 | 1 | 058 | 1 | 085 | 1 | 115 | 2 |
| 032 | 1 | 059 | 1 | 086 | 2 | 116 | 2 |
| 033 | 2 | 060 | 2 | 087 | 2 | | |

B

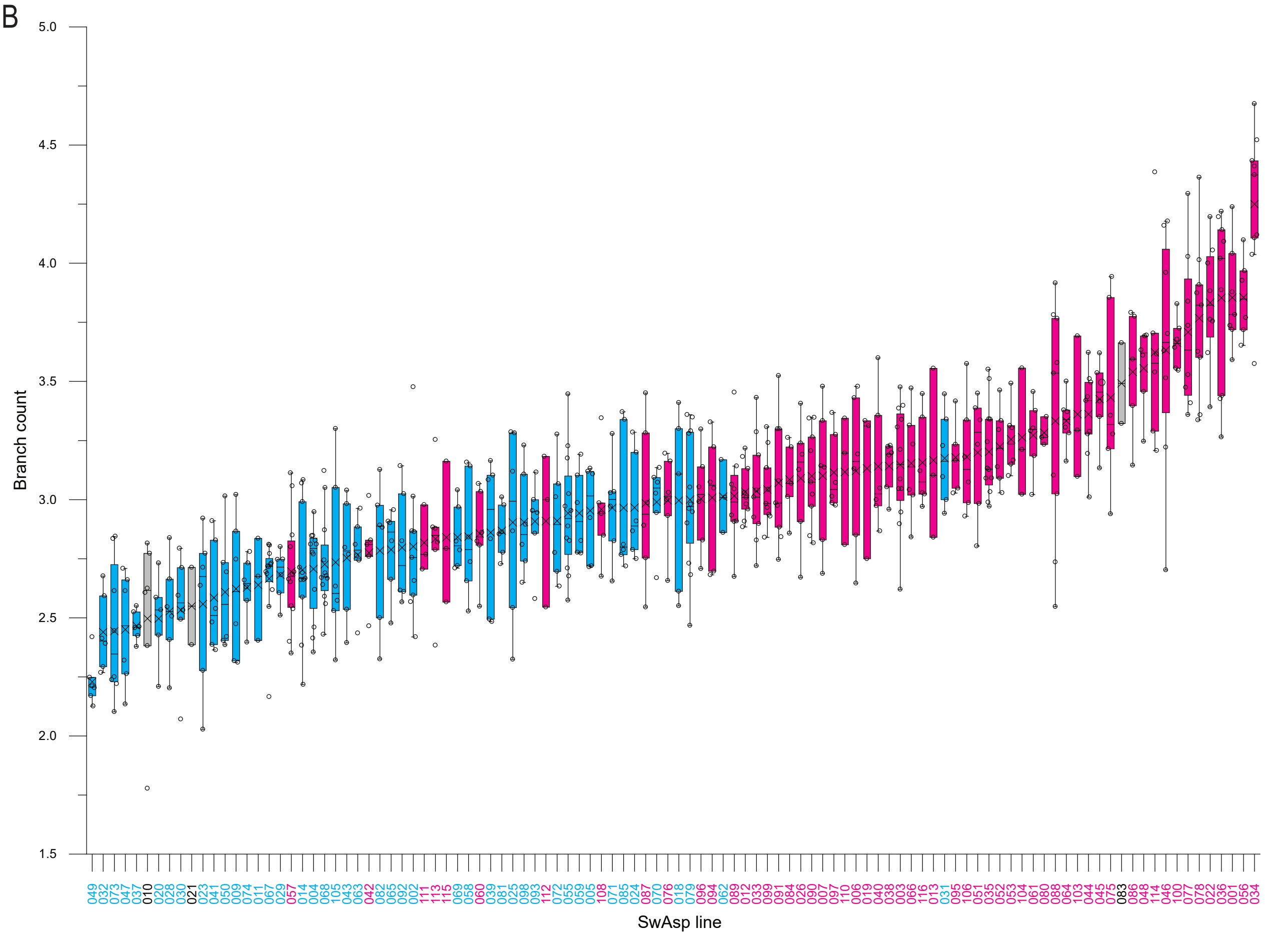


**Supplementary Fig. S14: *k*-means clustering assigns the Swedish aspen genotypes into more and less complex pavement cell shape clusters.** A, *k*-means cluster analysis of the SwAsp genotypes based on all cell shape and size features. The representative genotypes for more and less complex pavement cell shape are indicated in magenta and cyan, respectively. B, branch count of pavement cells in the SwAsp genotypes, as shown in Fig. 1D, but color-coded according to the cluster to which that genotype was assigned by *k*-means cluster analysis, with cyan used for cluster 1 and magenta for cluster 2.

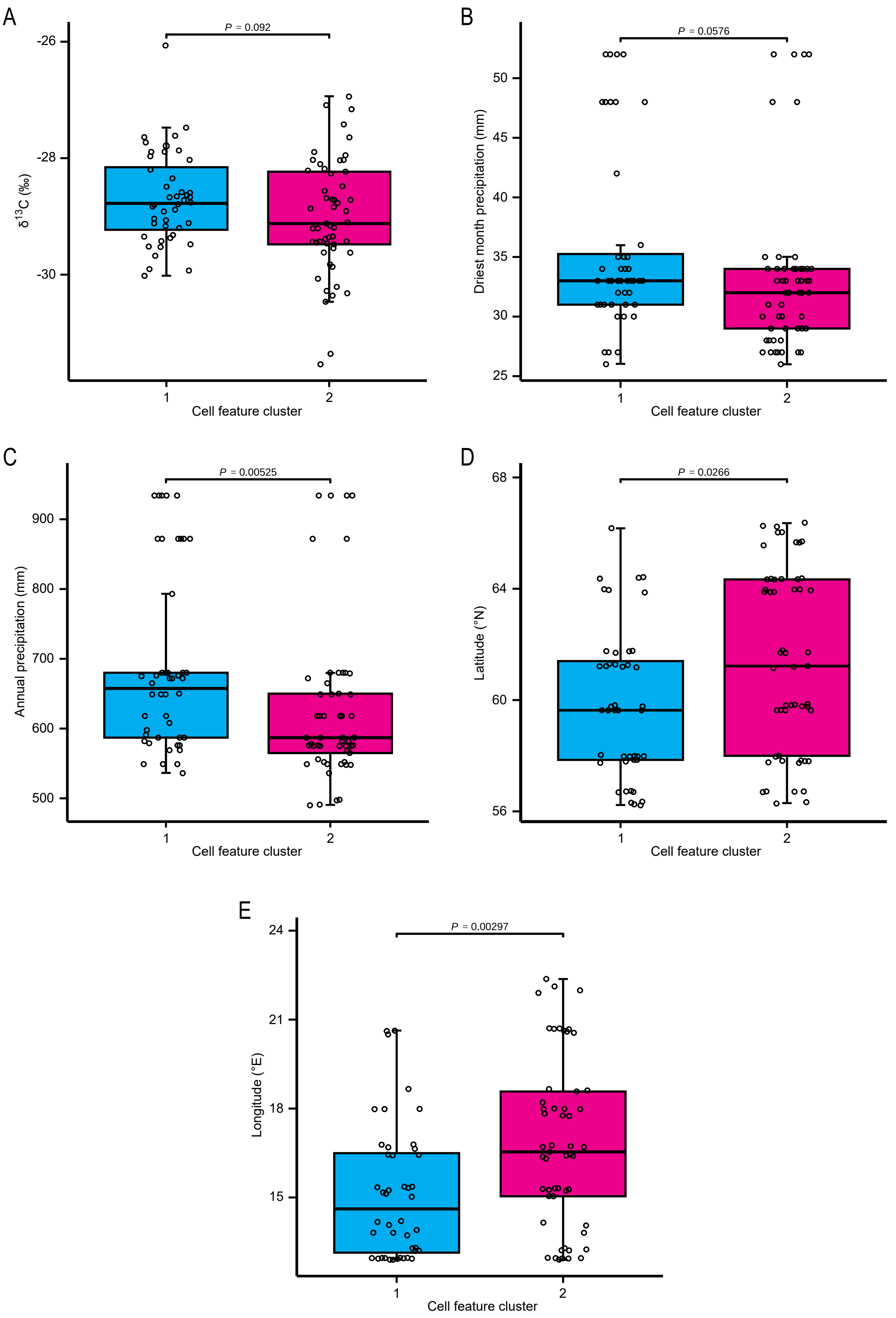


**Supplementary Fig. S15: The cell feature clusters associate significantly with original sampling site environments.** A-E, cluster assignment of the SwAsp genotypes from the *k*-means cluster analysis was used as a factor to partition the variance for foliar carbon isotope discrimination (delta$^{13}$C) (A), precipitation in the driest month of the year averaged over 30 years (B), annual precipitation averaged over 30 years (C), latitude (D) and longitude (E) at the original sampling sites. The *P*-values indicated were calculated using the Wilcoxon rank sum test.

**Supplementary Table S1: Primers used in this study.**

| Primer name | Primer sequence (5' to 3') |
|---|---|
| proPtMYB305a_F_attB1 | GGGGACAAGTTTGTACAAAAAAGCAGGCTTCGAAGGAGATAGAACCATGAATGGCGAATGGCAAATAAC |
| proPtMYB305a_ R_attB2 | GGGGACCACTTTGTACAAGAAAGCTGGGTGGTATAAAGGGAGGAATTAAGGAAAAG |
| PtMYB305a_CDS_F_attB1 | GGGGACAAGTTTGTACAAAAAAGCAGGCTTCGAAGGAGATAGAACCATGTCTTGGGGAGTGATGG |
| PtMYB305a_CDS_R_attB2 | GGGGACCACTTTGTACAAGAAAGCTGGGTGACAAAAAGGTGCCACTA |
| PtMYB305a_sgRNA1_T1F | ATATATGGTCTCGATTGTGGTCTGATCGAAGAGGGTGTTTTAGAGCTAGAAATAG |
| PtMYB305a_sgRNA2_T2R | ATTATTGGTCTCTAAACTCTCTTCAGGTCTGGTCTCCAATCTCTTAGTCGACTCTAC |
| PtMYB305a_sgRNA2_T2F | ATATTATTGGTCTCAAGATTGGAGACCAGACCTGAAGAGAGTTTTAGAGCTAGAAATAG |
| PtMYB305b_sgRNA1_T3F | ATATTATTGGTCTCAGTGATTGCAGGGCAGTTGCGCTGGGGGTTTTAGAGCTAGAAATAG |
| PtMYB305b_sgRNA3_T4R | ATTATTGGTCTCTAAACTGCCTTGTCTGAGTTGTCTCAATCACTACTTCGACTCTAG |
| PttMYB305a_2sgR_gF | CCCCCATATGCACAGAATTT |
| PttMYB305a_2sgR_gR | CAGTAAGGATGGTGGGAGTTC |
| PttMYB305b_4sgR_gF | AAGACAGCCTTTGCCAACAC |
| PttMYB305b_4sgR_gR | TTGCTGTTGTTGCTGCTGTA |
| PttMYB305a_RT-qPCR_F1 | TCTGATCGAAGAGGGTTGGA |
| PttMYB305a_RT-qPCR_R1 | GCCCTCTCTTCAGGTCTGGT |
| PttMYB305a_RT-qPCR_F2 | TGGCAATCTATGTGCAACAA |
| PttMYB305a_RT-qPCR_R2 | TTTGCGTTATCAAGGTTCCA |
| PtUBQ_RT-qPCR_F | GTTGATTTTTGCTGGGAAGC |
| PtUBQ_RT-qPCR_R | GATCTTGGCCTTCACGTTGT |
